# Multi-feature Classification to Improve Colorimetric Loop-Mediated Isothermal Amplification Fidelity

**DOI:** 10.64898/2026.06.03.728514

**Authors:** Gabrielle Melton, Daniel Antonio Negrón, Kelsey Hauser, Sveta Jagannathan, Nicholas Tolli, Katharine Jennings, Bryan D Necciai, Shanmuga Sozhamannan, Bradley W Abramson

## Abstract

Loop-mediated isothermal amplification (LAMP) is a cost-effective and portable assay technique for performing nucleic acid-based diagnostics in the field whose adoption is hindered by design and reproducibility issues. This is due to a complex primer design process that fine-tunes parameters across 6-8 binding regions. The likelihood of assay success depends on satisfying thermodynamic and secondary structure constraints while maintaining target specificity and avoiding overlaps between multiple primers. Software such as the <u>NEB® LAMP Primer Design Tool</u>, <u>PREMIER Biosoft LAMP Designer</u>, <u>Primer3</u>, <u>PCR Signature Erosion Tool (PSET)</u>, and <u>PrimerExplorer</u> enable automation of this task for researchers. However, in our experience, these programs can sometimes yield inconsistent results in laboratory testing. Here, we approached the issue by comparing and training multiple machine learning (ML) models on primer sets targeting various organisms from working assays and failing ones to determine significant features and improve predictions prior to ordering primer sets. A literature review produced an initial list of primer sets (n=116), which were then filtered down based on reference template availability to discern their FIP/BIP components (F2/F1c and B1c/B2). The final training set (n=109) included sequence and thermodynamic features derived from primers collected from the review (n=74) and those designed in-house with PSET (n=35). Failing assays were difficult to obtain from the publications, so we provided our own (n=23). Using WEKA Experimenter, models were created based on decision tree and Bayesian learning algorithms using an experimental scheme that performed a parameter grid search, seeded replicates, feature selection, and cross-validation while avoiding data-leakage and outputting logs for model comparison, feature analysis, and overfit assessment. Notably, thermodynamic features associated with the F1c and B1c primers consistently appeared in the top ranks according to consensus between information gain, class-correlation, and model-based feature ranking. For classification, the NaiveBayes algorithm had a TP and TN rate of 0.90 (± 0.02) and 0.73 (± 0.05) while achieving Cohen’s kappa coefficient and F-score values of 0.61 (± 0.06) and 0.91 (± 0.01). This work highlights how a practical model was built from a small, imbalanced training set incorporating negative research results, of which more are needed to improve generalization and refine parameters critical to assay success.

## Introduction

Loop-mediated isothermal amplification (LAMP) is a technique that offers a faster and cost-effective alternative to polymerase chain reaction (PCR) for field diagnostics^1^. LAMP has become increasingly common due to its simplicity, portability, pH-sensitive dyes, and because it bypasses the need for a thermocycler, while indicating amplification as an easily interpretable color change^2^. A disadvantage is that it requires 4-6 primers compared to the 2 and 3 for PCR and qPCR^3^. Multiple primer pairs enhance the specificity of amplification but also complicate the design and raise the risk of failure due to inadequate spacing and improper binding thermodynamics. Additional primers also increase the risk of deleterious secondary structure formation, such as heterodimers, homodimers, and hairpins that interfere with binding and amplification.

LAMP assays are practical for quickly performing pathogen surveillance, clinical diagnostics, allergen detection, environmental monitoring, veterinary surveillance, and antimicrobial resistance (AMR) analysis in the field, in austere environments, and in remote areas^4^. For example, they have been used to identify environmental contaminants and pathogens that threaten food and water safety, such as Shiga toxin-producing *E. coli* (STEC) and soybean allergens^5,6^. Recently, their use as a diagnostic tool has gained traction in global and public health initiatives. For example, organizations such as the WHO and CDC have employed LAMP to detect genes associated with AMR, such as those linked to methicillin-resistant *Staphylococcus aureus* (MRSA), providing a means of rapid detection for administering targeted antimicrobial therapies^7^. The multitude of LAMP applications highlights the importance of addressing primer design issues.

To design LAMP assays, researchers have created various software tools that enforce primer feature constraints. Programs such as Primer3, NEB LAMP Primer Design Tool, and PrimerExplorer, allow researchers to set parameter boundaries for various thermodynamic feature and sequence structure criteria^8^. Our group has also developed a LAMP design feature within our PCR Signature Erosion Tool (PSET) program^9^. Although primer sets are easily designed *in silico, in vitro* wet lab testing of some assays fail to amplify the target sequence or produce false positives, indicating that design tools may be ignoring critical feature optimizations. This can spiral into a time consuming and costly redesign and testing cycle.

Given the high degree of similarity of parameters from multiple design tools, we decided to narrow our focus to analyzing the thermodynamic features of primer sequences and their impact on overall assay success. Properties such as melting and annealing temperatures strongly influence primer-primer and primer-target interactions; therefore, we can calculate their contribution to overall assay success. Additional thermodynamic properties such as entropy, enthalpy, and Gibbs free energy reflect overall primer stability and can also predict the likelihood of secondary structure formation. Many primer design tools do not allow modifications to these thermodynamic properties during the design phase, apart from basic adjustments to melting temperature. Consequently, these parameters are potentially overlooked and unaccounted for in the primer design processes. We hypothesize that machine learning (ML) models can reveal parameters with the greatest influence on assay performance and predict success.

ML classification models can be used to determine feature importance within large datasets derived from PCR/qPCR primers^10,11^. For example, they can help understand complex primer design tasks or mis-amplification events^10,12,13^ or calculate confidence scores and likelihoods of assay success *in vitro*^14^. Some linear model classifiers for assay performance have been developed but generally focus on a single target region^15^. However, large and diverse datasets are required for ML methods to properly train models while avoiding bias and class imbalance issues.

In this study, we collected assays from the literature and included those developed in-house to build a training set based on the thermodynamic features of their primers. Models were built and optimized in a manner that mitigates overfit and avoids data-leakage. Decision tree and Bayesian algorithms were compared, with the NaiveBayes algorithm achieving the highest evaluation metrics, yielding a model that recovered the most true negatives. The ML scheme presented here highlights how even the few negative results we supplied for training can yield a practical model in the face of class imbalance, demonstrating the power of publishing failures.

## Materials and Methods

### Assay Design and Testing

LAMP assays were designed using PSET against an AMR pangenome from an earlier unpublished internal project and are indicated in Table S1 as “This study” under the source column. Primers were stored in 100-µM molecular grade water at −20°C. The LAMP reaction mix was prepared using 12.5-µL of WarmStart Colorimetric LAMP 2X Master Mix (New England Biolabs), 2.5-µL of reaction-specific LAMP 10X primer mix, and 8-µL molecular grade water. All LAMP reagents were thawed. Genomic DNA was isolated from *Enterobacter cloacae* complex (CDC AR isolate bank #AR-1150) by boil lysate method to provide a portion of the pangenome. Briefly, the sample was suspended in molecular grade water, heated to boil, and centrifuged. The resulting supernatant was retained as DNA isolate. Isolated DNA (1-µl, various concentrations) or molecular grade water (negative control) were loaded into each well of MicroAmp optical 96-well reaction plates (Applied Biosystems #N8010560). Reactions were performed in duplicate. Well contents were mixed by pipetting and incubated at 65°C for 30 minutes. Visual color determination of assay was used for successful or failed status.

### Feature Extraction

#### Assay Collection

LAMP assays were sourced from peer-reviewed literature that included the F3, B3, FIP, and BIP primers ^2,16–63^. The objective was to include a diverse set of assays covering a large set of organisms with a variety of primer configurations. All assays obtained from the literature were assumed to have working status. Those with failing status were determined *in vitro* based on those designed in-house and from a previous study of ours^16^. This data set is available in Table S1 with primer sequences listed as originally published. The F2/F1c and B1c/B2 primers were identified and extracted as described in the following sections or manually if possible.

#### Template Search

Prior to building the training set, the LAMP assays were filtered based on those whose primers were identified on a subject sequence with sufficient identity, orientation, and arrangement by analyzing the results from a BLAST+ (2.16.0+) run ^64^. The objective was to identify an ideal template for calculating some of the thermodynamic parameters as described in the following section and to also decompose the FIP/BIP primers into their F2/F1c and B1c/B2 constituents since not all publishers unambiguously listed them or demarcated their original boundaries (Table S1). The FASTA query file contained primer sequences with their linkers removed (“-TTTT-” or “-”) if present. Those containing ambiguous IUPAC DNA characters were expanded into all possible combinations as additional queries. Accordingly, the FASTA headers indicated the source assay, expansion number, and primer name to help with subsequent aggregation and filtering. BLAST+ was run with the “blastn-short” task to optimize for small sequences and with 100,000 alignments against a local copy of the NCBI Nucleotide “nt” database. The output format was adjusted to also report the subject strand. On a high-performance computing cluster, the query completed in 2:39:02 with 192 cores.

#### FIP/BIP Decomposition and Template Extraction

An R (v4.6.0) script was developed to process the BLAST+ results using the tidyverse (v2.0.0) package ^65^. BLAST+ hits were aggregated by assay id (parsed from the query header) and by subject accession. Percentage query coverage identity (PQCI) was computed for each hit as the percent identity multiplied by alignment length divided by query length. Subjects lacking a hit from a required primer were filtered out. A subsequent filter selected the top hits by PQCI, keeping the top F3/B3, (optional) LF/LB, and top two FIP/BIP hits, which were assumed to correspond to their F2/F1c and B1c/B2 components. Another filter checked for primer query order, strand, and overlap on each subject. Next, the FIP/BIP hits were checked for F2/F1c and B1c/B2 overlap and automatically adjusted if it was minimal (under 2-nt). Finally, FIP/BIP hits were filtered according to whether their hypothetical component lengths added up to their expected value. For each assay group, the subject sequence was demarcated with square brackets and parentheses corresponding to the identified primer and loop sequence hits using a fuzzy search algorithm to allow for minimal mismatches. An additional 6-nt were included up and downstream to provide additional binding context. Table S2 lists the filtered LAMP assay primer sets and target template.

#### Thermodynamics

A python (v3.13.13) script processed Table S2 to compute thermodynamic parameters using Pandas and Primer3 ^66,67^. Calculations included GC content, melting temperature, and thermodynamics related to hairpin, homodimer/heterodimer formation, and end stability. Heterodimer and end stability calculations were performed with respect to the identified template and its reverse complement. These calculations were also repeated for the last 5-nt and last 3-nt primer subsequences. For primers with ambiguous IUPAC letters, values were computed for each combinatorial expansion and averaged. A question mark indicated a missing value in the case of assays without loop primers. Table S3 includes the assay id, computed feature values, and nominal class label “success” (with “Y” for working/successful or “N” for failing status). Feature names contain underscores that delimit the calculation type (primary sequence “PS”, homodimer “HO”, heterodimer “HE”, and end stability “ES”), strand (agnostic/plus/minus “X”/”P”/ “N”), parameter (GC% “GC”, structure found “SF”, melting/annealing temperature “TM”, entropy “DS”, enthalpy “DH”, and Gibbs free energy “DG”), primer (“F3”, “F2”, “LF”, “F1c”, “B1c”, “B2”, “LB”, “B3”, “FIP”, and “BIP”), and sequence length (full length “FL”, last 5-nt “L5”, and last 3-nt “L3”). The script output a CSV, TSV, and ARFF (Attribute-Relation File Format) for direct use with the WEKA Java application (v3.9.7) ^68^.

### Machine Learning

#### Execution

A BASH script coordinated execution of WEKA on the command-line to calculate feature importance and ML model training. For variation and reproducibility, seeds for the internal pseudorandom number generator (PRNG) consisted of the first ten three-digit super-prime numbers (https://oeis.org/A006450). For ML, WEKA Experimenter was run within a loop cycling over the seeds and classifiers with corresponding model parameters. This and subsequent analysis scripts with corresponding output files are available in File S1.

#### Feature Ranking and Selection Stability

Features were ranked according to information gain (IG) via InfoGainAttributeEval/Ranker and correlation-based feature subset selection (CFS)^69^ via CfsSubsetEval/BestFirst with 10-fold cross-validation (CV) and a replicate with each of the ten PRNG seeds. Selection frequency for each feature was also calculated to determine persistence and consensus across non-baseline models based on how many times they were included in each cross-fold. Lustgarten’s stability measure (S) was also calculated for each model to compare pairwise feature selection stability across folds ^70^.

#### ML Model Training and Analysis

A hierarchical ML scheme was employed using WEKA Experimenter with 10-fold CV to run a MultiSearch (v2020.2.17) hyperparameter grid with 5-fold CV (instead of 10 to increase representation of the minority class). The former generated approximately 11 (⌈109/10⌉) instances of 8 (⌈83/10⌉) “Y” and 3 (⌈26/10⌉) “N” for testing and 98 (109 - 1⌉) instances with 75 (83 - 8) “Y” and 23 (26 - 3) “N” for MultiSearch tuning. This left the latter with approximately 20 (⌈98/5⌉) instances of 15 (⌈75/5⌉) “Y” and 5 (⌈23/5⌉) “N” for testing and 78 (98 - 20) instances with 60 (75 - 15) “Y” and (23 - 5) 18 “N” for training. MultiSearch wrapped a FilteredClassifier that performed attribute discretization via binning (to account for missing values from optional loop primers), IG selection (to immediately reduce high-noise features), and CFS (using GreedyStepwise with ranking) in this order within a MultiFilter to prevent data leakage. The final layer consisted of the classification algorithm, including three baseline (ZeroR, OneR, and DecisionStump) and four active learners. Of the latter, a simple model and complex model was included for the Bayesian (NaiveBayes and BayesNet) and tree-based (J48 and RandomForest) methods respectively ^71,72^. For all classifiers, MultiSearch generated grid values for the “numToSelect” parameter of the GreedyStepwise filter to enforce a hard limit for the number of features to incorporate, ranging from 1 to 20. Two additional search axes were included for parameterizable classifiers with the first and second exploring an additional 3 and 4 values, thereby maintaining an equal grid size for fair downstream analysis. Grid axes are described in Table S4. This workflow was repeated for each PRNG seed. With 8 cores, the workflow ran in 17:40.43. Optimal model parameters were determined by calculating the top per-fold model configuration based on Cohen’s kappa coefficient (κ) and then keeping the modal value of each individual parameter. Model performance was assessed via Tukey’s range test (α=0.05) on observed κ values across folds and seeds using the agricolae (v1.3-7) R package ^73^.

## Results

### Assay Testing

Only our assays were verified to have working or failing status *in vitro*, with the rest from the literature outside of our group assumed to work. LAMP primer sets were validated *in vitro* using lysate from target DNA from an *E. cloacae* complex CDC AR isolate (Figure S1). Of the 19 tested primer sets, one had a true positive result indicating assay success. For the remaining primer sets, 15 had true negative results, two had false positive results, and one had an inconclusive result. True negatives, inconclusive results, and false positives were classified as assay failures, resulting in a total of 18 primer sets with failing status. This status is reflected as a “Y” or “N” for success or failure, respectively, under the “success” class column in Table S2. Edited and raw photos of the LAMP results are available in File S1.

### Template Search and Training Set

The BLAST+ query identified suitable templates for 74 of the 116 assays obtained from the peer-reviewed literature for the R script to extract the F2/F1c and B1c/B2 primers. Combined with our in-house assays, the python script calculated thermodynamic values from the primer sequences and corresponding templates, yielding a training set with 109 instances and 640 features in Table S3. For loop primers, 56 assays had both, 3 had LF only, 29 had LB only, and 21 had none.

### Feature Performance

Figure 1 compares the top 30 features ranked by the IG (A), CFS (B), and model consensus-based methods corresponding to the data in Table S5. The first two were evaluated on the entire training set to determine the most informative features based on individual and collective/nonredundant strength respectively. These two methods selected the same 19 features deriving from at least one of each non-FIP/BIP primer sequence (F3x2, F2x2, F1cx3, LFx2, LBx2, B1cx5, B2x1, and B3x2). The model consensus heatmap (C) shows how often each feature was selected within each cross-fold. All methods ranked the same 8 features within the top 30 and were primarily comprised of those derived from the B1c (n=5) and F1c (n=3) primers. Only primary sequence TM, heterodimer, and hairpin features were found in common across the methods.

**Figure 1.**
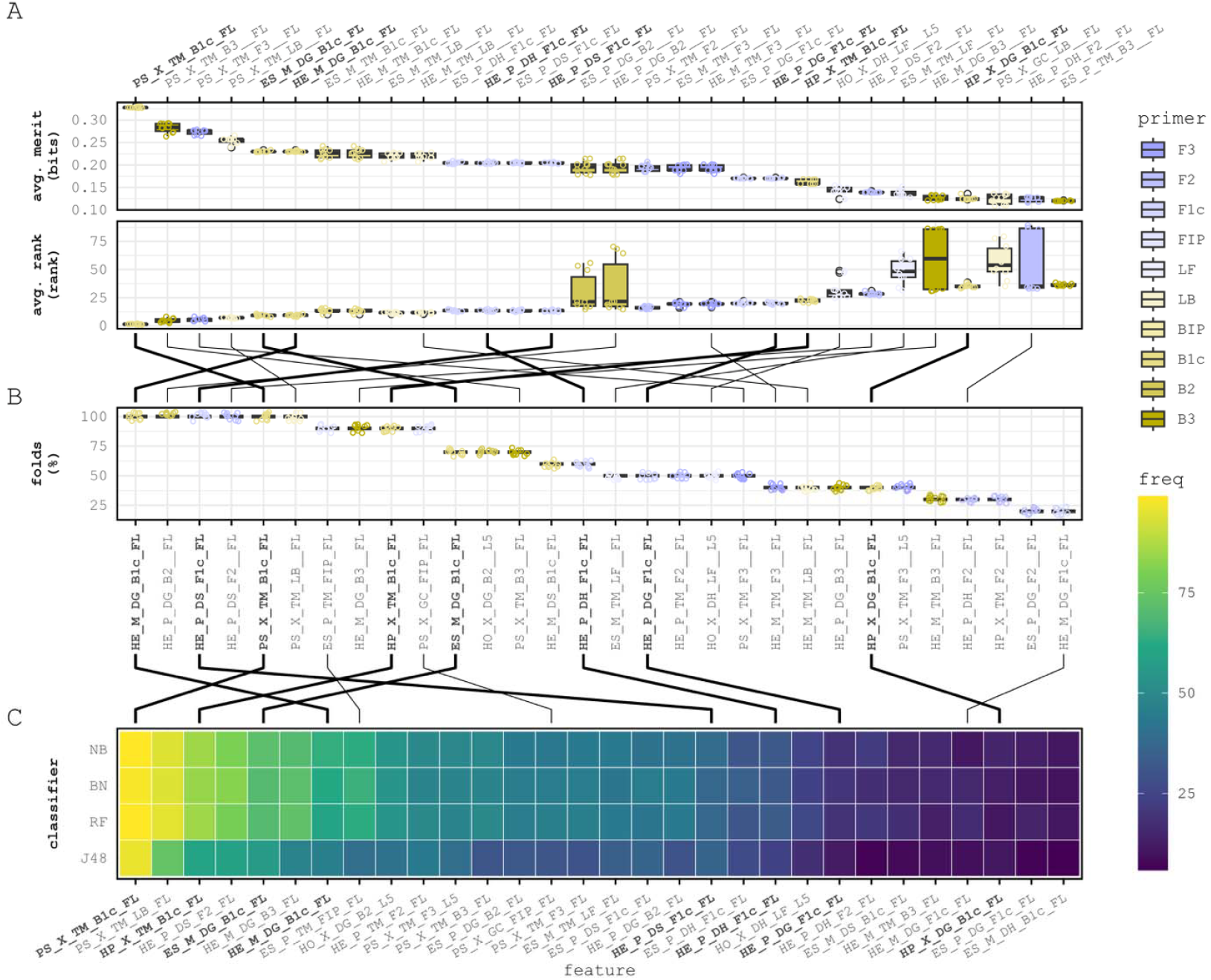
Feature ranking. A, B, and C correspond to InfoGainAttributeEval+Ranker and CfsSubsetEval+BestFirst ranking distributions and model-based selection frequency. Lines indicate the change in rank of shared features between each method. Bold lines highlight the features selected across all three methods, which were derived from the B1c and F1c. Underscores delimit feature properties: calculation (primary sequence “PS”, homodimer “HO”, heterodimer “HE”, and end stability “ES”), strand (agnostic “X” and plus/minus “P”/ “N”), sequence/thermodynamic property, primer, and sequence length (full length “FL”, last 5-nt “L5”, and last 3-nt “L3”).

### Model Performance

Table 1 and Table S6 list the model performance metrics broken down by class. All models outperformed baseline classifiers with respect to κ, except for J48. Models were grouped by CLD (compact letter display) based on Tukey’s range test (α=0.05) (Table S7), resulting in four distinct tiers. Accordingly, members of the top performing group (CLD=a) had moderate-to-substantial^74^ κ values and are therefore indistinguishable by this value alone. However, despite similar confusion matrix values, NaïveBayes notably achieved a much higher true positive rate for class N. The failure of J48 and underperformance of RandomForest are likely due to the lack of minority class values providing sufficient tree splitting information and instances overall for tree depth.

**Table 1.** Model performance metrics. Models are grouped by CLD according to Tukey’s range test (α=0.05). The κ and S columns are Cohen’s Kappa coefficient and Lustgarten’s stability measure. All cells contain average and standard deviation values.

| model | CLD | $\kappa$ | S | AUROC | Class Y (n=83) | | | Class N (n=26) | | |
| --- | --- | --- | --- | --- | --- | --- | --- | --- | --- | --- |
|  |  |  |  |  | TPR | PPV | F-score | TPR | PPV | F-score |
| Naïve Bayes | a | 0.61<br>± 0.06 | 0.59<br>± 0.13 | 0.90<br>± 0.03 | 0.90<br>± 0.02 | 0.92<br>± 0.01 | 0.91<br>± 0.01 | 0.73<br>± 0.05 | 0.70<br>± 0.05 | 0.72<br>± 0.04 |
| Bayes Net | a | 0.57<br>± 0.10 | 0.60<br>± 0.14 | 0.89<br>± 0.03 | 0.91<br>± 0.02 | 0.90<br>± 0.02 | 0.90<br>± 0.02 | 0.66<br>± 0.09 | 0.71<br>± 0.08 | 0.68<br>± 0.08 |
| Random Forest | a | 0.55<br>± 0.05 | 0.59<br>± 0.13 | 0.89<br>± 0.04 | 0.93<br>± 0.02 | 0.89<br>± 0.02 | 0.90<br>± 0.01 | 0.61<br>± 0.06 | 0.73<br>± 0.04 | 0.66<br>± 0.04 |
| OneR | b | 0.39<br>± 0.08 | 0.64<br>± 0.21 | 0.69<br>± 0.03 | 0.92<br>± 0.02 | 0.85<br>± 0.01 | 0.88<br>± 0.02 | 0.46<br>± 0.06 | 0.63<br>± 0.08 | 0.53<br>± 0.06 |
| J48 | bc | 0.38<br>± 0.15 | 0.60<br>± 0.21 | 0.76<br>± 0.07 | 0.89<br>± 0.03 | 0.85<br>± 0.04 | 0.87<br>± 0.03 | 0.49<br>± 0.13 | 0.58<br>± 0.12 | 0.53<br>± 0.12 |
| Decision Stump | c | 0.29<br>± 0.09 | 0.70<br>± 0.35 | 0.67<br>± 0.04 | 0.83<br>± 0.03 | 0.85<br>± 0.02 | 0.82<br>± 0.03 | 0.48<br>± 0.07 | 0.48<br>± 0.07 | 0.48<br>± 0.06 |
| ZeroR | d | 0.00<br>± 0.00 | 0.56<br>± 0.09 | 0.50<br>± 0.00 | 1.00<br>± 0.00 | 0.76<br>± 0.00 | 0.86<br>± 0.00 | 0.00<br>± 0.00 | NaN<br>± NA | NaN<br>± NA |

Figure 2 compares model performance and feature selection stability. The A plot shows the observed κ values across every fold and seed of each classifier. B traces the average κ observed over the grid search axis of the “numToSelect” parameter, which controls the number of model features. Circles highlight the value resulting in the highest observed average κ from the set of best parameter combinations, with values peaking within the grid search range. The C plot shows the observed Lustgarten’s S (Table S7) between the selected feature sets of pairwise folds for each classifier and D compares the average value to the corresponding average κ of each model. Optimal model parameters are available in Table S9.

**Figure 2.**
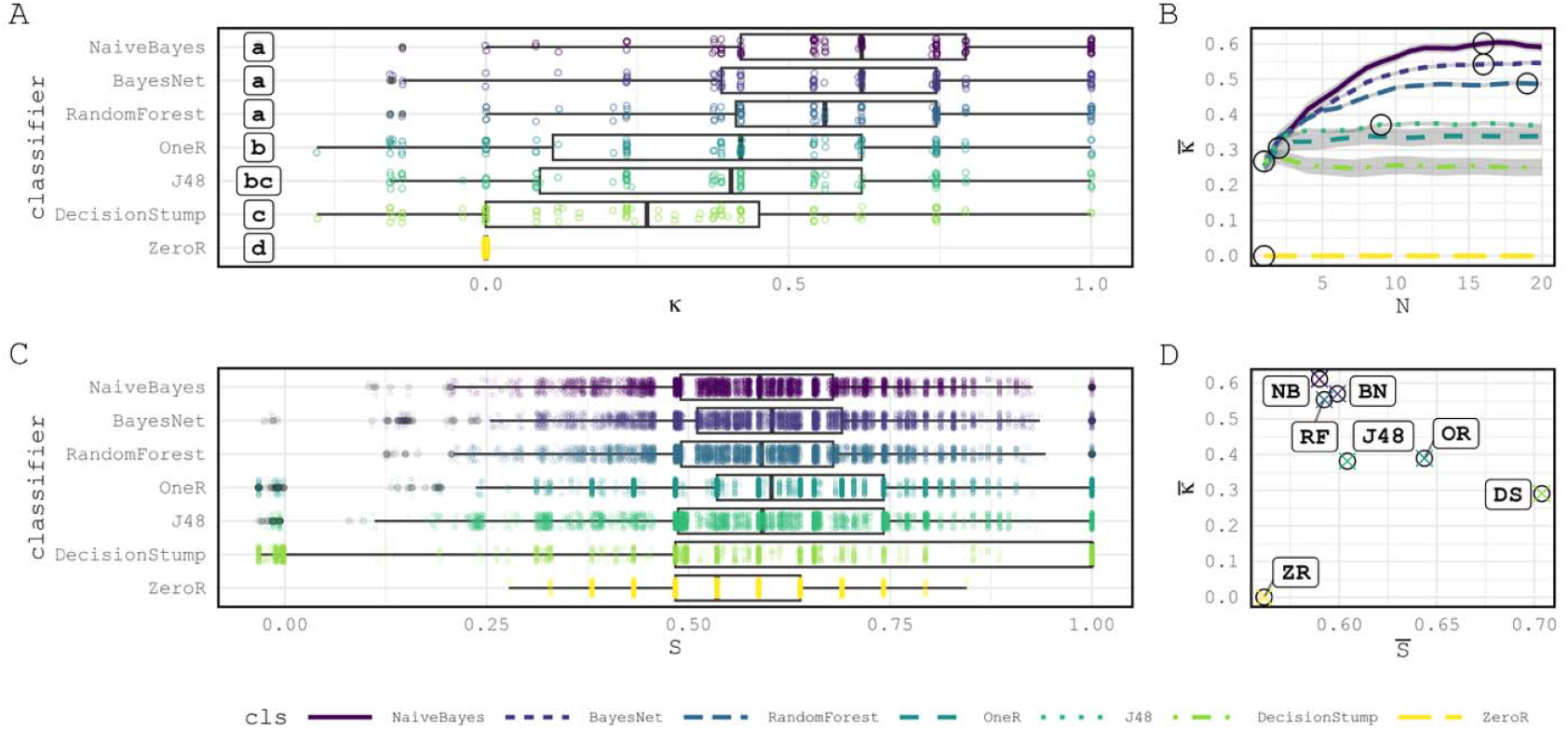
Model comparison and feature selection stability. The box plots in A correspond to the Cohen’s Kappa coefficient (κ) values observed for each seed and cross-fold for each classifier, which are grouped by Tukey’s range test CLD labels (α=0.05). B traces the optimal number of features selected over the grid search for that parameter. C plots the distribution of Lustgarten’s metric (S) values between pairwise folds across seeds for each classifier. D plots the average κ and S observed for each model (with initials indicated).

## Discussion

Here we present a diverse, curated collection of LAMP assays and an ML scheme for optimizing feature selection and classification. Published LAMP assays (n=116) were utilized and filtered by template availability (n=74) alongside newly designed ones (n=35), some of which were successful in laboratory testing or failed to reliably amplify the target sequence. Template target sequences were not recovered for all assays collected from the literature review. This could be due to only having performed one BLAST+ search with one set of parameters or that the original researchers used sequences outside of GenBank for their designs. Additionally, the R script FIP/BIP primer decomposition algorithm may have used overly strict thresholds when assessing subject hits.

The LAMP primer sets were deconstructed to calculate thermodynamic parameters for inclusion into the training set. Features such as melting temperature, entropy, enthalpy, and Gibbs free energy have a large influence on how the primers bind, dimerize, form hairpins, or successfully amplify target DNA regions on the template. For example, undesirable secondary structures should exhibit a low melting temperature and near-zero Gibbs free energy, indicating that they are unstable and unlikely to form. Accordingly, the python script ran thermodynamic analysis functions from Primer3 to build the training set. The great majority were uninformative or noisy according to initial feature analysis. This was due to calculating the same features from different lengths of the same primers, increasing the number of redundant or useless features.

The consensus set of features across ranking methods included those associated with the F1c and B1c primers. In the top ranks, the F2 and B2 were also present and shared between the IG and CFS methods. This is notable since they are themselves derived from the FIP and BIP primers, which are responsible for forming the dumbbell structures required for amplification^2^. Therefore, the models benefitted from the FIP/BIP decomposition script presented here. Accordingly, the full FIP/BIP sequence features did not rank highly as their constituent primers were more informative. Thermodynamic features associated with these primers may represent those parameters most critical to assay success and potentially require further scrutiny when designing LAMP assay sets.

Models of varying complexity were included in the ML scheme derived from decision and probabilistic algorithms. For example, J48 builds a single tree whereas RandomForest builds an ensemble. Similarly, the simpler NaiveBayes classifiers assumes feature independence compared to BayesNet, which looks for dependencies. The idea was to vary model interpretability, parsimony, and ability to handle correlated features and small data sets. Accordingly, NaiveBayes excelled in exploiting the distribution, bias, and variance within this small data set. Similarly, the small training set and class imbalance may have significantly impacted the tree-based methods. Increasing training set instances with failing assays may ultimately reveal BayesNet, RandomForest, or another model as the stronger classifier, especially as it would provide more information for stronger rankings and conditional probabilities. Future directions should explore the inclusion of additional features, such as non-specific amplification (NSA) probabilities of secondary structures or ideal binding ^14^. Synthetic data generation via SMOTE^75^ or cost-based classification with CostSensitiveClassifier^68^ may also alleviate class-imbalance issues.

The ML scheme presented here successfully built a predictive model that exceeded evaluation benchmarks and baseline classifiers, showing that there is a useful signal from a small, unbalanced training set. However, the training set included failing assays designed from only from the PSET program. Therefore, it is unknown how it would generalize to predict failures originating from other design tools, examples of which are virtually unavailable from the literature. This is a common trend in published research, in which negative results and their associated assay information such as primer sequences are generally not reported/published or considered meaningful for impactful publications. As AI/ML are integrated into every aspect of life, the models will be limited by published datasets of only successful assays, which is a huge detriment to each scientific field and should be reconsidered as agentic AI is becoming available to parse published datasets. By incorporating all tested materials into a publication, including failed tests, additional insights may be discovered. The lack of information from failed experiments/assays in the public record is sure to limit future ML studies, such as this one, and researchers should consider reporting their beautiful failures in future publications.

The advanced capabilities of ML models are particularly relevant to LAMP assay primer design, where the requirement of four to six primers introduces a broader array of features and variables for analysis. Manual and *in vitro* wet lab evaluation of such extensive data sets is not only impractical but also inadequate for grasping the intricate relationships between features and their impacts on assay success in the laboratory. Deploying classifiers into the design cycle shows promise as they can provide immediate predictive results prior to primer synthesis, enhancing efficiency, and minimizing the time required for troubleshooting when assays are later tested in the laboratory. This may help mitigate failures and significantly improve public health response by optimizing primer design to target a wide array of DNA targets, thereby improving diagnostic research related to viral and pathogen detection and antimicrobial resistance.

## Author Contributions

Conceptualization: GM, DN, SS, & BA. Methodology: GM, DN, KH, NT, & BA. Software: GM, DN, & BA. Validation: GM, DN, SJ, NT, & BA. Formal analysis: GM, DN, NT, & BA. Investigation: GM, DN, KH, SJ, NT, & BA. Resources: GM, KH, NT, & BA. Data Curation: GM, DN, NT, & BA. Writing - Original Draft: GM, DN, & BA. Writing - Review & Editing: GM, DN, KH, SJ, NT, KJ, BN, SS, & BA. Visualization: DN & BA. Supervision: DN, NT, & BA. Project administration: KJ & BA. Funding acquisition: BN, SS, & BA.

## Acknowledgments

The views expressed in this manuscript are those of the authors and do not necessarily reflect the official policy or position of the CPE-CBRND, the Department of War, or the U.S. Government. This work was prepared as part of the author(s) official duties. Title 17 U.S.C. § 105 provides that ‘Copyright protection under this title is not available for any work of the United States Government.’ Title 17 U.S.C. §101 defines U.S.

Government work as work prepared by a military service member or employee of the U.S. Government as part of that person’s official duties. References to non-federal entities or their products do not constitute or imply Department of War’s or Army’s endorsement of any company, product or organization. Funding for this study was provided and executed by the Capability Program Executive for Chemical, Biological, Radiological and Nuclear Defense’s (CPE-CBRND) Joint Project Lead for CBRN Enabling Technologies (JPL CBRN ET) on behalf of the Department of War’s Chemical and Biological Defense Program. Thank you to Dr. Arturo Donate for reviewing.

## Supplementary Material

Supplementary Table S1: LAMP assays included in this study.

Supplementary Table S2: LAMP primer sets with decomposed FIP/BIP sequences.

Supplementary Table S3: Training set of computed thermodynamic values.

Supplementary Table S4: Parameter grid.

Supplementary Table S5: Feature rankings.

Supplementary Table S6: Confusion matrix and model statistics.

Supplementary Table S7: Tukey’s range test.

Supplementary Table S8: Lustgarten’s stability measure.

Supplementary Table S9: Optimal model parameters.

Supplementary Table SA: Matrix of author contributions by CRediT Taxonomy.

Supplementary Figure S1: LAMP assay photo and description (within File S1).

Supplementary File S1: Zip file containing code, data, figures, and photos.

