## Supplemental code and data. for "Multi-feature Classification to Improve Colorimetric Loop-Mediated Isothermal Amplification Fidelity": featurize.py

```
#!/usr/bin/env python

from collections import namedtuple
from itertools import chain, product
from operator import attrgetter

import pandas as pd
from primer3.bindings import *
from primer3.p3helpers import reverse_complement as rc
from pset.assay import Assay

amb = dict(zip(
    ("A", "C", "G", "T", "R", "Y", "S", "W", "K", "M", "B", "D", "H", "V", "N"),
    ("A", "C", "G", "T", "AG", "CT", "GC", "AT", "GT", "AC", "CGT", "AGT", "ACT", "ACG", "ACGT")
))

thermo_fields = "structure_found", "tm", "ds", "dh", "dg"
thermo_hdr = "SF", "TM", "DS", "DH", "DG"
primers = "F3", "F2", "LF", "F1c", "B1c", "B2", "LB", "B3"

def expand(seq):
    yield from ("".join(ele) for ele in product(*(amb[ele] for ele in seq)))


def gc(seq):
    return sum(ele in "GC" for ele in seq) / len(seq)


def avg_thermo(results):
    return namedtuple("ThermoResult", thermo_fields)(*(sum(ele)/len(ele) for ele in zip(*map(attrgetter(*thermo_fields), results))))


def main():
    # Load data
    path = "supplement.xlsx"
    df_data = pd.read_excel(path, sheet_name="S2")

    # Calculate features
    assays = []
    results = []
    slicers = {"FL": lambda x: x, "L5": lambda x: x[-5:]}
    for record in df_data.to_records():
        assay = Assay.factory(record.definition)
        assays.append((record.id, *(assay.components.get(key, "") for key in primers)))
        for slice_mode, slice_func in slicers.items():
            for primer, seq in chain(assay.components.items(), zip(("FIP", "BIP"), (assay.fip(), assay.bip()))):
                seq = seq if primer in {"F3", "F2", "B1c", "LB"} else rc(seq)
                seqs = list(expand(slice_func(seq)))
                defn_fwd = "".join(ele for ele in assay.definition if ele not in "[]()")
                defn_rev = rc(defn_fwd)
                results.append(
                    [
                        record.id,
                        slice_mode,
                        primer,
                        sum(map(gc, seqs)) / len(seqs),  # GC content
                        sum(map(calc_tm, seqs)) / len(seqs),  # Melting temperature
                        *(avg_thermo(map(calc_hairpin, seqs))),  # Hairpin formation
                        *(avg_thermo(map(calc_homodimer, seqs))),  # Homodimer formation
                        *(avg_thermo((calc_heterodimer(seq, defn_fwd) for seq in seqs))),  # Heterodimer formation +/+
                        *(avg_thermo((calc_heterodimer(seq, defn_rev) for seq in seqs))),  # Heterodimer formation +/-
                        *(avg_thermo((calc_end_stability(seq, defn_fwd) for seq in seqs))),  # End stability +/+
                        *(avg_thermo((calc_end_stability(seq, defn_rev) for seq in seqs))),  # End stability +/-
                    ]
                )

    # output primers
    df = pd.DataFrame(assays, columns=("id", *primers))
    df.to_csv("S2.csv", index=False)
    df.to_csv("S2.tsv", index=False, sep="\t")

    # output training set
    columns = [
        "id", "length", "primer",
        "PS_X_GC", "PS_X_TM",
        *(f"{p}_{q}" for p, q in product(("HP_X", "HO_X", "HE_P", "HE_M", "ES_P", "ES_M"), thermo_hdr))
    ]
    df = pd.DataFrame(pd.pivot(pd.DataFrame(results, columns=columns), index="id", columns=["primer", "length"]).to_records()).fillna("?")
    df.columns = [df.columns[0], *(ele[2:-2].replace("', '", "_") for ele in df.columns[1:])]
    df = df.merge(df_data[["id", "success"]], how="left", on="id")
    ## as CSV/TSV
    df.to_csv("S3.csv", index=False)
    df.to_csv("S3.tsv", index=False, sep="\t")
    ## as ARFF
    with open("S3.arff", "w") as file:
        print(
            "@RELATION LAMP",
            *(f"@ATTRIBUTE {ele} NUMERIC" for ele in df.columns[1:-1]),
            f"@ATTRIBUTE {df.columns[-1]} {{{','.join(df[df.columns[-1]].unique())}}}",
            "@DATA",
            sep="\n",
            file=file
        )
    with open("S3.arff", "a") as file:
        df.drop(columns=["id"]).to_csv(file, header=False, index=False)

    return 0


if __name__ == "__main__":
    main()
```
