## Supplemental code and data. for "Multi-feature Classification to Improve Colorimetric Loop-Mediated Isothermal Amplification Fidelity": filter1.R

```
library(tidyverse)

# make an id representing the primer set loop configuration
loop.id <-
  openxlsx2::read_xlsx("supplement.xlsx", sheet = "S1") %>%
  with(setNames(1 + if_else(is.na(LF), 0, 1) + 2 * if_else(is.na(LB), 0, 1), id))

# primer configurations
primers <- list(c("F3", "B3", "FIP", "BIP"), c("F3", "B3", "FIP", "BIP", "LF"), c("F3", "B3", "FIP", "BIP", "LB"), c("F3", "B3", "FIP", "BIP", "LF", "LB"))
# fwd primer id orders
pcheck1 <- c("F3FIPFIPBIPBIPB3", "F3FIPLFFIPBIPBIPB3", "F3FIPFIPBIPLBBIPB3", "F3FIPLFFIPBIPLBBIPB3")
# rev primer id orders
pcheck2 <- c("B3BIPBIPFIPFIPF3", "B3BIPBIPFIPLFFIPF3", "B3BIPLBBIPFIPFIPLF", "B3BIPLBBIPFIPLFFIPF3")
# strand configurations
scheck <- c("++-+--", "++--+--", "++-++--", "++--++--")

# load primers
df.primer <-
  openxlsx2::read_xlsx("supplement.xlsx", sheet = "S1") %>%
  mutate(id = as.character(id)) %>%
  remove_rownames() %>%
  filter(!(source %in% c("doi:10.1186/s12859-024-06001-3", "This study"))) %>%
  select(id, matches("^([FB](3|2|1|IP)c?|L[FB])$", ignore.case = F)) %>%
  pivot_longer(-id, names_to = "primer", values_to = "seq", values_drop_na = T) %>%
  mutate(len = str_length(seq))

# read/process BLAST+ hits
df.hit1 <-
  read_tsv(
    "blast-hits-subset.tsv",
    comment = "#",
    col_names = c("qaccver", "saccver", "idn", "alen", "mis", "gap", "qstart", "qend", "sstart", "send", "evalue", "score", "strand"),
    col_types = "ffdiiiiiiiddc"
  ) %>%
  # parse-out query header info: assay id, ambiguous expansion, primer id
  separate_wider_delim(qaccver, delim = "-", names = c("id", "exp", "primer")) %>%
  select(-exp) %>%
  filter(str_detect(primer, "^([FB](3|IP)?|L[FB])$")) %>%
  mutate(
    strand = if_else(strand == "plus", "+", "-"),
    start = if_else(sstart < send, sstart, send),
    end = if_else(sstart < send, send, sstart)
  ) %>%
  select(-sstart, -send) %>%
  left_join(select(df.primer, id, primer, len), by = join_by(id, primer)) %>%
  mutate(
    pqcov = idn * alen / len,
    loop = loop.id[id],
    nprimer = sapply(primers[loop], length)
  ) %>%
  rename(qlen = len) %>%
  group_by(id, saccver) %>%
  filter(n_distinct(primer) == nprimer)

# keep the top hits to each subject for each asay id
# only the top of each F3/B3/LF/LB
# top two FIP/BIP
df.hit2 <-
  bind_rows(
    df.hit1 %>%
      filter(primer %in% c("F3", "B3", "LF", "LB")) %>%
      group_by(id, saccver, primer) %>%
      slice_max(n = 1, order_by = pqcov, with_ties = F),
    df.hit1 %>%
      filter(primer == "FIP") %>%
      group_by(id, saccver, primer) %>%
      slice_max(n = 2, order_by = pqcov, with_ties = F),
    df.hit1 %>%
      filter(primer == "BIP") %>%
      group_by(id, saccver, primer) %>%
      slice_max(n = 2, order_by = pqcov, with_ties = F)
  )

# check primer and strand configuration and overlap violations
df.result <-
  df.hit2 %>%
  group_by(id, saccver) %>%
  group_modify(\(.x, .y) {
    .x <- arrange(.x, start)
    idx <- loop.id[[.y$id]]
    pc <- ifelse(.x$primer[1] == "F3", pcheck1[idx], pcheck2[idx])
    sc <- scheck[idx]
    p.ok <- paste0(.x$primer, collapse = "") == pc
    s.ok <- paste0(.x$strand, collapse = "") == sc
    o.ok <- !any(.x$end < lead(.x$start, 0))
    if (p.ok && s.ok && o.ok) {
      .x
    } else {
      data.frame()
    }
  })

# adjust FIP/BIP overlaps <= 2-nt and extend if necessary
# keep those whose components sum to FIP/BIP length
df.result2 <-
  df.result %>%
  arrange(primer, qstart, .by_group = T) %>%
  group_modify(\(.x, .y) {
    df.IP <-
      filter(.x, endsWith(primer, "IP")) %>%
      mutate(primer = paste(primer, c(1, 2), sep = "-"))
    # overlap
    df.IP[[2, "qstart"]] <- with(df.IP, ifelse(abs(qend[1] - qstart[2]) <= 2, qend[1] + 1, qstart[2]))
    df.IP[[4, "qstart"]] <- with(df.IP, ifelse(abs(qend[3] - qstart[4]) <= 2, qend[3] + 1, qstart[4]))
    df.IP[[2, "qend"]] <- with(df.IP, ifelse((qlen[2] - qend[2]) <= 1, qlen[2], qend[2]))
    df.IP[[4, "qend"]] <- with(df.IP, ifelse((qlen[4] - qend[4]) <= 1, qlen[4], qend[4]))
    # extend
    ok.BIP <-
      df.IP[[1, "qend"]] - df.IP[[1, "qstart"]] + 1 + df.IP[[2, "qend"]] - df.IP[[2, "qstart"]] + 1 ==
        df.IP[[2, "qlen"]]
    ok.FIP <-
      df.IP[[3, "qend"]] - df.IP[[3, "qstart"]] + 1 + df.IP[[4, "qend"]] - df.IP[[4, "qstart"]] + 1 ==
        df.IP[[4, "qlen"]]
    # filter
    if (ok.BIP && ok.FIP) {
      bind_rows(df.IP, filter(.x, !endsWith(primer, "IP")))
    } else {
      data.frame()
    }
  })

# keep subjects with FIP/BIP components with full coverage from start to length
# keep those subjects with all required components
df.result3 <-
  df.result2 %>%
  filter(
    str_detect(primer, "[FB]IP-2$") | qstart == 1,
    str_detect(primer, "[FB]IP-1$") | qend == qlen
  ) %>%
  mutate(group = cur_group_id()) %>%
  group_by(id, saccver) %>%
  group_modify(\(.x, .y) {
    if (length(setdiff(primers[[.x$loop[1]]], str_remove(.x$primer, "-[12]$"))) == 0) {
      .x
    } else {
      data.frame()
    }
  }) %>%
  group_by(id) %>%
  slice_min(group, n = 1, with_ties = T) %>%
  arrange(id, qstart)

# write subject accession ranges to file
df.result3 %>%
  summarise(
    saccver = first(saccver),
    start = max(1, min(start) - 6 - 1),
    end = max(end) + 6 + 1
  ) %>%
  arrange(id) %>%
  select(-id) %>%
  write_tsv("blast-saccver-subset.tsv", col_names = F)

# write metadata for second phase
df.result3 %>%
  arrange(id, start) %>%
  mutate(primer = str_remove(primer, "-[12]$")) %>%
  left_join(df.primer, join_by(id, primer)) %>%
  ungroup() %>%
  mutate(
    pstart = pmin(qstart, qend),
    pend = pmax(qstart, qend),
    pseq = if_else(endsWith(primer, "IP"), str_sub(seq, pstart, pend), seq),
    pseq = str_replace(pseq, "-TTTT-", "-"),
    pseq = str_remove_all(pseq, "-"),
    idx = cumsum(id != lag(id, 1, default = "0"))
  ) %>%
  group_by(idx) %>%
  saveRDS("filter.rds")
```
