## Supplemental code and data. for "Multi-feature Classification to Improve Colorimetric Loop-Mediated Isothermal Amplification Fidelity": filter2.R

```
library(tidyverse)

# IUPAC ambiguous letters and reverse complement
dna <-setNames(strsplit("TGCAAWSKMYRVHDBN", "")[[1]], strsplit("ACGTUWSMKRYBDHVN", "")[[1]])
# reverse complement
rc <- \(seq) paste0(dna[rev(strsplit(seq, "")[[1]])], collapse = "")
# IUPAC ambiguous letters mapping
amb <-
  setNames(
    sapply(
      c("A", "C", "G", "T", "AG", "CT", "GC", "AT", "GT", "AC", "CGT", "AGT", "ACT", "ACG", "ACGT"),
      str_split,
      "",
      USE.NAMES = F
    ),
    c("A", "C", "G", "T", "R", "Y", "S", "W", "K", "M", "B", "D", "H", "V", "N")
  )
# create seq with regex for all ambiguous letters
ambreg <-
  function(seq) {
    strsplit(seq, "")[[1]] %>%
      amb[.] %>%
      sapply(\(e) ifelse(nchar(e <- paste0(e, collapse = "")) == 1, e, str_glue("[{e}]"))) %>%
      paste0(collapse = "")
  }
# legal brackets
brackets <- c("[][][][][][]", "[][]()[][][][]", "[][][][]()[][]", "[][]()[][]()[][]")

# load primers
df.primer <- 
  openxlsx2::read_xlsx("supplement.xlsx", sheet = "S1") %>%
  mutate(id = as.character(id)) %>%
  remove_rownames() %>%
  filter(!(source %in% c("doi:10.1186/s12859-024-06001-3", "This study"))) %>%
  select(id, matches("^([FB](3|2|1|IP)c?|L[FB])$", ignore.case = F)) %>%
  pivot_longer(-id, names_to = "primer", values_to = "seq", values_drop_na = T) %>%
  mutate(len = str_length(seq))

# load subjects
saccver <-
  read_lines("blast-saccver-subset.fna") %>%
  as_tibble() %>%
  mutate(group = cumsum(startsWith(value, ">"))) %>%
  group_by(group) %>%
  summarise(seq = paste0(tail(value, n = -1), collapse = "")) %>%
  pull(seq)

readRDS("filter.rds") %>%
  group_map(\(.x, .y) {
    id <- .x$id[1]
    sseq <- saccver[.y$idx]
    sseq <- ifelse(.x$primer[1] == "F3", sseq, rc(sseq))
    if (.x$primer[1] == "B3") {
      .x <- map_df(.x, rev)
    }
    keys <- c("F2", "F1c")
    FIP <- df.primer[df.primer$id == id & df.primer$primer %in% keys, ]$seq
    if (length(FIP) < 2 | any(nchar(FIP) < 15)) {
      FIP <- c(.x[.x$primer == "FIP", ][[1, "pseq"]], .x[.x$primer == "FIP", ][[2, "pseq"]])
    }
    keys <- c("B1c", "B2")
    BIP <- df.primer[df.primer$id == id & df.primer$primer %in% keys, ]$seq
    if (length(BIP) < 2 | any(nchar(BIP) < 15)) {
      BIP <- c(.x[.x$primer == "BIP", ][[1, "pseq"]], .x[.x$primer == "BIP", ][[2, "pseq"]])
    }
    qseqs <-
      c(
        .x[.x$primer == "F3", ][[1, "pseq"]],
        .x[.x$primer == "B3", ][[1, "pseq"]],
        .x[.x$primer == "FIP", ][[1, "pseq"]], .x[.x$primer == "FIP", ][[2, "pseq"]],
        .x[.x$primer == "BIP", ][[1, "pseq"]], .x[.x$primer == "BIP", ][[2, "pseq"]],
        ifelse("LF" %in% .x$primer, .x[.x$primer == "LF", ][[1, "pseq"]], ""),
        ifelse("LB" %in% .x$primer, .x[.x$primer == "LB", ][[1, "pseq"]], "")
      ) %>%
      Filter(nchar, .)
    # gotta use a for-loop due to loop dependency
    for (idx in seq_along(qseqs)) {
      areg <- aregexec(ambreg(qseqs[idx]), sseq)
      if (!is.na(areg) & areg[1] == -1) {
        qseqs[idx] <- rc(qseqs[idx])
        areg <- aregexec(ambreg(qseqs[idx]), sseq)
      }
      pattern <- str_sub(sseq, areg[[1]], areg[[1]] + attr(areg[[1]], "match.length") - 1)
      sseq <- 
        ifelse(
          areg[1] >= 1 & !str_detect(pattern, "[\\[\\]\\(\\)]"),
          str_replace(
            sseq,
            pattern,
            ifelse(idx <= 6, str_glue("[{qseqs[idx]}]"), str_glue("({qseqs[idx]})"))
          ),
          NA
        )
    }
    with(.x, data.frame(id = id[1], saccver = saccver[1], start = min(start), stop = max(end), definition = sseq)
    )
  }) %>%
  bind_rows() %>%
  filter(str_remove_all(definition, "[^\\[\\]\\(\\)]") %in% brackets) %>%
  mutate(definition = str_extract(definition, "([A-Z]{0,6}\\[.*][A-Z]{0,6})")) %>%
  filter(complete.cases(.)) %>%
  write_tsv("blast-definition-subset.tsv")
```
