## Supplemental code and data. for "Multi-feature Classification to Improve Colorimetric Loop-Mediated Isothermal Amplification Fidelity": plots.R

```
library(tidyverse)
library(patchwork)

read_multisearch <- function(path) {
  lines <- read_lines(path)
  params <-
    lines %>%
    Filter(\(ele) str_detect(ele, "^ - [0-9]+\\. dimension:"), .) %>%
    str_extract("^ - [0-9]+\\. dimension: ([^,]+),.*", group = T) %>%
    unique() %>%
    paste0("p.", .)
  lines %>%
    as_tibble() %>%
    mutate(
      value = str_squish(value),
      fold = cumsum(value == "=== Initial space - Start ===")
    ) %>%
    filter(str_detect(value, "^Performance \\(.*\\): [0-9]+\\.?[0-9]* \\(.*\\): ")) %>%
    separate_wider_delim(value, delim = ": ", names = c("perf", "value", "cached")) %>%
    mutate(perf = str_sub(str_remove(perf, "Performance "), 2, -2)) %>%
    separate_wider_delim(perf, delim = ", ", names = params) %>%
    mutate(
      cls = first(Filter(\(ele) startsWith(ele, "Classifier:"), lines)),
      cls = str_split_i(cls, ": ", -1),
      stat = str_extract(value, "\\((.*)\\)", group = T),
      value = str_split_i(value, " ", 1),
      path = path,
      across(all_of(c(params, "value")), as.numeric)
    ) %>%
    select(cls, fold, everything())
}

read_eval_cfs <- function(path) {
  lines <- read_lines("data/WEKA-CfsSubsetEval-109.out.txt")
  idx <- which(lines == "number of folds (%)  attribute")
  lines[(idx + 1):(length(lines) - 1)] %>%
    str_squish() %>%
    str_replace_all("[(%)]", " ") %>%
    str_replace_all(" \\s+", " ") %>%
    as_tibble() %>%
    separate_wider_delim(value, delim = " ", names = c("nfolds", "pfolds", "attr_idx", "attr_key")) %>%
    mutate(
      nfolds = as.integer(nfolds),
      pfolds = as.numeric(pfolds),
      attr_idx = as.integer(attr_idx),
      path = path
    )
}

read_eval_info <- function(path) {
  lines <- read_lines(path)
  idx <- which(lines == "average merit      average rank  attribute")
  lines[(idx + 1):(length(lines) - 1)] %>%
    str_squish() %>%
    str_replace_all("\\+-", " ") %>%
    str_replace_all("\\s+", " ") %>%
    as_tibble() %>%
    separate_wider_delim(
      value,
      delim = " ",
      names = c("avg_merit", "avg_merit_sd", "avg_rank", "avg_rank_sd", "attr_idx", "attr_key")
    ) %>%
    mutate(
      across(starts_with("avg_", ignore.case = F), as.numeric),
      attr_idx = as.integer(attr_idx),
      path = path
    )
}

read_zip_log <- function(log) {
  df.meta <-
    archive::archive(log) %>%
    mutate(
      seed = str_extract(path, "^([0-9]+)\\.", group = T),
      fold = str_extract(path, "^[0-9]+\\.([0-9]+)\\.", group = T),
      cls = str_extract(path, "-W_([^_]+)(?!.*-W)", group = T),
      cls = str_extract(cls, "\\.(.*)", group = T),
      cls = str_remove_all(cls, "\\(.*")
    )
  bind_cols(
    df.meta,
    bind_rows(
      lapply(1:nrow(df.meta), \(ele) {
        lines <- read_lines(archive::archive_read(log, ele))
        bind_cols(
          data.frame(
            stat = str_split(grep("Evaluation: ", lines, value = T), "[:, ]")[[1]][3]
          ),
          as_tibble(as.list(setNames(
            Filter(str_length, tail(str_split(grep("^Coordinates: ", lines, value = T), "[:, ]")[[1]], n = -2)),
            paste0("p.", str_split_i(grep("^[0-9]+\\. parameter: property: ", lines, value = T), "[:, ]", 6))
          ))),
          as_tibble(as.list(
            str_split_i(head(grep("^@attribute ", lines, value = T), n = -1), " ", 2) %>%
              {
                setNames(rep(1, length(.)), paste0("a.", .))
              }
          ))
        )
      })
    ) %>%
      select(stat, everything())
  )
}

ls.cls.baseline <- c("ZeroR", "OneR", "DecisionStump")
ls.metric <-
  setNames(
    c(
      "Num_true_positives",
      "Num_false_negatives",
      "Num_true_negatives",
      "Num_false_positives",
      "True_positive_rate",
      "False_negative_rate",
      "True_negative_rate",
      "False_positive_rate",
      "Kappa_statistic",
      "IR_precision",
      "F_measure",
      "Matthews_correlation_coefficient",
      "Area_under_ROC",
      "Area_under_PRC",
      "Mean_absolute_error",
      "Root_mean_squared_error",
      "Relative_absolute_error",
      "Root_relative_squared_error"
    ),
    c(
      "TP", "FN", "TN", "FP", "TPR", "FNR", "TNR", "FPR",
      "κ", "PPV", "F-score", "MCC", "AUROC", "AUPRC",
      "MAE", "RMSE", "RAE", "RRSE"
    )
  )
ls.primers <- c("F3", "F2", "F1c", "FIP", "LF", "LB", "BIP", "B1c", "B2", "B3")
ls.attr.names <- c("calc", "strand", "prop", "primer", "length")

df.csv <-
  list.files("data", pattern = "^WEKA-[^-]+-[0-9]{3}\\.out\\.csv$", full.names = T) %>%
  parallel::mclapply(read_csv, col_types = cols(.default = "c"), mc.cores = 8) %>%
  bind_rows() %>%
  rename(cls = Key_Scheme_options) %>%
  mutate(
    cls = str_extract(cls, "-W ([^ ]+)(?!.*-W )", group = T),
    cls = str_split_i(cls, "\\.", -1),
    cls = str_remove_all(cls, "\\W"),
    ruf = str_glue("{Key_Run}_{Key_Fold}"),
    across(any_of(unname(ls.metric)), as.numeric),
    across(c(ruf, cls), as.factor)
  )

df.log <-
  list.files("data", pattern = "^WEKA-[^-]+-[0-9]{3}\\.log\\.txt$", full.names = T) %>%
  parallel::mclapply(read_multisearch, mc.cores = 8) %>%
  bind_rows() %>%
  mutate(
    cls = str_extract(cls, "-W ([^ ]+)", group = T),
    cls = str_split_i(cls, "\\.", -1),
    seed = str_split_i(basename(path), "[\\.\\-]", 3)
  ) %>%
  rename_with(~ gsub(".*\\.", "p.", .), starts_with("p.", ignore.case = F)) %>%
  select(cls, seed, fold, starts_with("p.", ignore.case = F), everything())

df.eva.cfs <-
  list.files("data", pattern = "^WEKA-CfsSubsetEval+-[0-9]{3}\\.out\\.txt$", full.names = T) %>%
  parallel::mclapply(read_eval_cfs) %>%
  bind_rows() %>%
  separate_wider_delim(attr_key, delim = "_", names = ls.attr.names, cols_remove = F) %>%
  mutate(primer = if_else(nchar(primer) == 3, primer, paste0(primer, "_"))) %>%
  unite(attr_key, all_of(ls.attr.names), remove = F) %>%
  mutate(primer = factor(str_remove(primer, "_"), levels = ls.primers))

df.eva.nfo <-
  list.files("data", pattern = "^WEKA-InfoGainAttributeEval+-[0-9]{3}\\.out\\.txt$", full.names = T) %>%
  parallel::mclapply(read_eval_info) %>%
  bind_rows() %>%
  separate_wider_delim(attr_key, delim = "_", names = ls.attr.names, cols_remove = F) %>%
  mutate(primer = if_else(nchar(primer) == 3, primer, paste0(primer, "_"))) %>%
  unite(attr_key, all_of(ls.attr.names), remove = F) %>%
  mutate(primer = factor(str_remove(primer, "_"), levels = ls.primers))

df.zip <-
  list.files("data", pattern = "^WEKA-[^-]+-[0-9]{3}\\.out\\.zip$", full.names = T) %>%
  parallel::mclapply(read_zip_log, mc.cores = 8) %>%
  bind_rows() %>%
  select(cls, starts_with("a.", ignore.case = F))

df.csv.con <-
  df.csv %>%
  group_by(Key_Run, cls) %>%
  summarize(
    across(any_of(Filter(\(ele) startsWith(ele, "Num"), ls.metric)), sum),
    across(any_of(Filter(\(ele) !startsWith(ele, "Num"), ls.metric)), mean),
    TPR.N = TNR,
    FNR.N = FPR,
    TNR.N = TPR,
    FPR.N = FNR,
    PPV.N = TN / (TN + FN),
    .groups = "drop"
  ) %>%
  group_by(cls) %>%
  # TODO: output for supplement
  summarize(
    across(
      -one_of("Key_Run"),
      ~ sprintf("%0.2f (± %.2f)", mean(.x), sd(.x))
    )
  ) %>%
  rename_with(~ paste0(.x, ".Y"), .cols = matches("[CRVe]$", ignore.case = F))

df.csv.tuk <-
  df.csv %>%
  lm(Kappa_statistic ~ cls + ruf, data = .) %>%
  agricolae::HSD.test("cls", group = T) %>%
  {
    left_join(
      rownames_to_column(.$groups, "cls"),
      rownames_to_column(.$means, "cls"),
      by = join_by(cls, Kappa_statistic)
    )
  } %>%
  rename(CLD = groups)

df.eva.cfs.top <-
  df.eva.cfs %>%
  group_by(attr_key) %>%
  summarize(pfolds = mean(pfolds)) %>%
  slice_max(pfolds, n = 30, with_ties = F)

df.eva.nfo.top <-
  df.eva.nfo %>%
  group_by(attr_key) %>%
  summarize(avg_merit = mean(avg_merit)) %>%
  slice_max(avg_merit, n = 30, with_ties = F)

df.log.mod <-
  df.log %>%
  select(cls, seed, fold, starts_with("p.", ignore.case = F), value) %>%
  arrange(across(starts_with("p.", ignore.case = F))) %>%
  group_by(cls, seed, fold) %>%
  slice_max(value, with_ties = F) %>%
  ungroup() %>%
  select(-seed, -fold) %>%
  group_by(across(all_of(starts_with("p.", ignore.case = F)))) %>%
  rename(kappa = value) %>%
  pivot_longer(-c(cls, kappa), values_drop_na = T) %>%
  group_by(cls, name, value) %>%
  summarize(n = n(), av = mean(kappa), .groups = "drop") %>%
  arrange(value) %>%
  group_by(cls, name) %>%
  slice_max(n, with_ties = F) %>%
  ungroup() %>%
  arrange(cls) %>%
  mutate(
    cls = factor(cls, levels = df.csv.tuk$cls),
    name = factor(name, levels = Filter(\(ele) startsWith(ele, "p."), names(df.log)))
  ) %>%
  arrange(cls, name)

num.features <- length(unique(df.eva.nfo$attr_key))
df.zip.sta <-
  df.zip %>%
  mutate(across(everything(), ~ replace_na(.x, 0))) %>%
  group_by(cls) %>%
  group_modify(\(.x, .y) {
    x <-
      as.matrix(.x) %>%
      tcrossprod() %>%
      {
        cbind(
          .[lower.tri(.)],
          t(combn(diag(.), m = 2))
        )
      }
    colnames(x) <- c("r", "k1", "k2")
    x %>%
      as_tibble(.name_repair = "check_unique") %>%
      mutate(
        E = k1 * k2 / num.features,
        S = (r - E) / (pmin(k1, k2) - E)
      )
  }) %>%
  mutate(cls = factor(cls, levels = rev(df.csv.tuk$cls)))

df.zip.con <-
  df.zip %>%
  filter(!(cls %in% ls.cls.baseline)) %>%
  pivot_longer(-cls, values_drop_na = T) %>%
  group_by(cls, name) %>%
  summarize(freq = sum(value), .groups = "drop") %>%
  complete(cls, name, fill = list(freq = 0)) %>%
  arrange(cls, freq) %>%
  {
    filter(., name %in% slice_max(filter(., cls == df.csv.tuk$cls[1]), freq, n = 30, with_ties = F)$name)
  } %>%
  mutate(
    cls = factor(cls, levels = rev(df.csv.tuk$cls)),
    name = str_remove_all(name, "^a\\."),
    name = reorder(name, freq, .fun = sum, decreasing = T)
  ) %>%
  rename(attr_key = name)

df.rnk.1 <-
  data.frame(
    p = df.eva.nfo.top$attr_key,
    q = df.eva.cfs.top$attr_key
  ) %>%
  rowid_to_column(var = "group") %>%
  pivot_longer(-group, values_to = "x") %>%
  mutate(
    y = (seq_along(x) %% 2),
    x = factor(x, levels = df.eva.nfo.top$attr_key)
  ) %>%
  group_by(group) %>%
  group_modify(\(.x, .y) {
    if (any(is.na(.x$x))) {
      mutate(.x, x = first(x), y = first(y))
    } else {
      f <- if (as.numeric(first(.x$x)) < as.numeric(last(.x$x))) desc else identity
      bind_rows(.x, mutate(.x, y = y + 0.25)) %>%
        arrange(f(y))
    }
  }) %>%
  rename(attr_key = x)

df.rnk.2 <-
  data.frame(
    p = df.eva.cfs.top$attr_key,
    q = levels(df.zip.con$attr_key)
  ) %>%
  rowid_to_column(var = "group") %>%
  pivot_longer(-group, values_to = "x") %>%
  mutate(
    y = (seq_along(x) %% 2),
    x = factor(x, levels = df.eva.cfs.top$attr_key)
  ) %>%
  group_by(group) %>%
  group_modify(\(.x, .y) {
    if (any(is.na(.x$x))) {
      mutate(.x, x = first(x), y = first(y))
    } else {
      f <- if (as.numeric(first(.x$x)) < as.numeric(last(.x$x))) desc else identity
      bind_rows(.x, mutate(.x, y = y + 0.25)) %>%
        arrange(f(y))
    }
  }) %>%
  rename(attr_key = x)

ls.rnk.all <-
  bind_rows(df.rnk.1, df.rnk.2) %>%
  ungroup() %>%
  count(attr_key) %>%
  filter(n == 8) %>%
  pull(attr_key)

ls.rnk.1.grp <-
  df.rnk.1 %>%
  arrange(desc(y)) %>%
  filter(row_number() == n(), attr_key %in% ls.rnk.all) %>%
  pull(group)

ls.rnk.2.grp <-
  filter(df.rnk.2, any(attr_key %in% ls.rnk.all), n() == 4) %>%
  pull(group) %>%
  unique()

p1 <-
  df.eva.nfo %>%
  select(attr_key, primer, avg_merit, avg_rank) %>%
  filter(attr_key %in% df.eva.nfo.top$attr_key) %>%
  pivot_longer(-c(attr_key, primer)) %>%
  mutate(attr_key = factor(attr_key, levels = df.eva.nfo.top$attr_key)) %>%
  ggplot(aes(attr_key, value)) +
  geom_boxplot(aes(fill = primer), linewidth = 0.5, outlier.shape = 1, show.legend = T) +
  geom_jitter(aes(color = primer), width = 0.25, pch = 1, size = 1, show.legend = F) +
  facet_grid(
    name ~ .,
    labeller = as_labeller(c(avg_merit = "avg. merit\n(bits)", avg_rank = "avg. rank\n(rank)")),
    scales = "free_y",
    switch = "y"
  ) +
  scale_x_discrete(position = "top") +
  scale_color_discrete(drop = F, palette = "Blue-Yellow 3") +
  scale_fill_discrete(drop = F, palette = "Blue-Yellow 3") +
  labs(x = NULL, y = NULL, tag = "A") +
  theme_minimal() +
  theme(
    axis.text.x = element_text(
      angle = 30, 
      hjust = 0, 
      vjust = 0, 
      size = 8, 
      face = sapply(df.eva.nfo.top$attr_key, \(ele) ifelse(ele %in% ls.rnk.all, "bold", "plain"))
    ),
    axis.ticks.x = element_line(color = "black"),
    axis.title = element_text(size = 10),
    panel.border = element_rect(color = "black", linewidth = 1),
    plot.margin = unit(c(0, 5.5, 0, 5.5), "pt"),
    strip.placement = "outside",
    text = element_text(family = "mono")
  )

p2 <-
  df.rnk.1 %>%
  mutate(linewidth = if_else(group %in% ls.rnk.1.grp, 0.75, 0.25)) %>%
  ggplot(aes(attr_key, y, group = group)) +
  geom_line(aes(linewidth = linewidth), color = "black") +
  facet_grid("" ~ ., scales = "free_y") +
  scale_linewidth_identity() +
  scale_y_reverse() +
  labs(x = NULL, y = NULL) +
  theme_minimal() +
  theme(
    axis.text = element_blank(),
    panel.border = element_blank(),
    panel.grid = element_blank(),
    plot.margin = unit(c(0, 5.5, 0, 5.5), "pt"),
    text = element_text(family = "mono")
  )

p3 <-
  df.eva.cfs %>%
  filter(attr_key %in% df.eva.cfs.top$attr_key) %>%
  mutate(attr_key = factor(attr_key, levels = df.eva.cfs.top$attr_key)) %>%
  ggplot(aes(attr_key, pfolds)) +
  geom_boxplot(aes(fill = primer), linewidth = 0.5, outlier.shape = 1, show.legend = T) +
  geom_jitter(aes(color = primer), width = 0.25, pch = 1, size = 1, show.legend = F) +
  facet_grid("folds\n(%)" ~ ., switch = "y") +
  scale_color_discrete(drop = F, palette = "Blue-Yellow 3") +
  scale_fill_discrete(drop = F, palette = "Blue-Yellow 3") +
  labs(x = NULL, y = NULL, tag = "B") +
  theme_minimal() +
  theme(
    axis.text.x = element_text(
      angle = 90, 
      hjust = 1, 
      vjust = 0.5, 
      size = 8, 
      face = sapply(df.eva.cfs.top$attr_key, \(ele) ifelse(ele %in% ls.rnk.all, "bold", "plain"))
    ),
    axis.ticks.x = element_line(color = "black"),
    axis.title = element_text(size = 10),
    legend.position = "none",
    palette.colour.discrete = "Blue-Yellow 3",
    panel.border = element_rect(color = "black", linewidth = 1),
    plot.margin = unit(c(0, 5.5, 0, 5.5), "pt"),
    strip.placement = "outside",
    text = element_text(family = "mono")
  )

p4 <-
  df.rnk.2 %>%
  mutate(linewidth = if_else(group %in% ls.rnk.2.grp, 0.75, 0.25)) %>%
  ggplot(aes(attr_key, y, group = group)) +
  geom_line(aes(linewidth = linewidth), color = "black") +
  facet_grid("" ~ ., scales = "free_y") +
  scale_linewidth_identity() +
  scale_y_reverse() +
  labs(x = NULL, y = NULL) +
  theme_minimal() +
  theme(
    axis.text = element_blank(),
    panel.border = element_blank(),
    panel.grid = element_blank(),
    plot.margin = unit(c(0, 5.5, 0, 5.5), "pt"),
    text = element_text(family = "mono")
  )

p5 <-
  df.zip.con %>%
  ggplot(., aes(x = attr_key, y = cls, fill = freq)) +
  geom_tile(color = "white") +
  facet_grid("classifier" ~ ., scales = "free_y", switch = "y") +
  scale_y_discrete(labels = with(df.csv.tuk, setNames(str_remove_all(df.csv.tuk$cls, "[^0-9A-Z]"), cls))) +
  scale_fill_viridis_c() +
  labs(x = "feature", y = NULL, tag = "C") +
  theme_minimal() +
  theme(
    axis.text.x = element_text(
      angle = 30, 
      hjust = 1, 
      vjust = 1, 
      size = 8, 
      face = sapply(levels(df.zip.con$attr_key), \(ele) ifelse(ele %in% ls.rnk.all, "bold", "plain"))
    ),
    axis.ticks.x = element_line(color = "black"),
    axis.title = element_text(size = 10),
    legend.key.height = unit(1, "null"),
    panel.border = element_rect(color = "black", linewidth = 1),
    plot.margin = unit(c(0, 5.5, 0, 5.5), "pt"),
    strip.placement = "outside",
    text = element_text(family = "mono")
  )

p1 / p2 / plot_spacer() / p3 / p4 / plot_spacer() / p5 +
  plot_layout(heights = c(3, 1, -0.5125, 1.5, 1, -0.5125, 2), guides = "collect") &
  theme(
    plot.tag = element_text(size = 16, family = "mono"),
    strip.text = element_text(face = "bold")
  )
ggsave("Fig1.pdf", dpi = 300)
ggsave("Fig1.png", dpi = 300)

p6 <-
  df.csv %>%
  left_join(select(df.csv.tuk, cls, CLD), by = join_by(cls)) %>%
  mutate(cls = factor(cls, levels = df.csv.tuk$cls)) %>%
  ggplot(aes(Kappa_statistic, cls)) +
  geom_boxplot(outlier.size = 1, outlier.alpha = 0.5) +
  geom_jitter(aes(, color = cls), pch = 1, size = 1, alpha = 0.5, height = 0.25) +
  geom_label(
    df.csv.tuk,
    mapping = aes(label = CLD),
    x = min(df.csv$Kappa_statistic),
    family = "mono",
    fontface = "bold",
    alpha = 0.5
  ) +
  scale_color_viridis_d() +
  scale_y_discrete(limits = rev) +
  guides(color = "none") +
  theme_minimal() +
  labs(x = expression(kappa), y = "classifier") +
  theme(
    legend.position = "none",
    panel.border = element_rect(color = "black", linewidth = 1),
    text = element_text(family = "mono")
  )

p7 <-
  df.zip.sta %>%
  mutate(cls = factor(cls, levels = df.csv.tuk$cls)) %>%
  ggplot(aes(x = S, y = cls)) +
  geom_boxplot(outlier.size = 1, outlier.alpha = 0.1) +
  geom_jitter(aes(color = cls), height = 0.25, pch = 1, alpha = 0.1, size = 0.5) +
  scale_color_viridis_d() +
  scale_y_discrete(limits = rev) +
  guides(color = "none") +
  labs(y = "classifier") +
  theme_minimal() +
  theme(
    legend.position = "none",
    panel.border = element_rect(color = "black", linewidth = 1),
    text = element_text(family = "mono")
  )

p8 <-
  df.log %>%
  mutate(cls = factor(cls, levels = df.csv.tuk$cls)) %>%
  group_by(cls, p.numToSelect) %>%
  summarise(
    av = mean(value),
    sd = sd(value),
    se = sd / sqrt(n()),
    lo = av - qt(0.975, df = n() - 1) * se,
    up = av + qt(0.975, df = n() - 1) * se,
    .groups = "drop"
  ) %>%
  {
    ggplot(., aes(p.numToSelect, av, group = cls)) +
      geom_ribbon(aes(ymin = lo, ymax = up), alpha = 0.25) +
      geom_line(aes(linetype = cls, color = cls), linewidth = 1) +
      geom_point(
        data = left_join(
          pivot_wider(filter(df.log.mod, name == "p.numToSelect")),
          .,
          by = join_by(cls, p.numToSelect),
          suffix = c(".x", "")
        ),
        mapping = aes(x = p.numToSelect, y = av),
        size = 4,
        pch = 1
      ) +
      guides(
        linetype = guide_legend(nrow = 1),
        color = guide_legend(nrow = 1)
      ) +
      scale_y_continuous(breaks = seq(0, 1, .1)) +
      scale_color_viridis_d() +
      labs(x = "N", y = expression(bar(kappa))) +
      theme_minimal() +
      theme(
        legend.key.width = unit(0.5, "in"),
        legend.text = element_text(size = 7),
        panel.border = element_rect(color = "black", linewidth = 1),
        text = element_text(family = "mono")
      )
  }

p9 <-
  summarize(df.zip.sta, S = mean(S)) %>%
  left_join(
    summarize(group_by(df.csv, cls), kappa = mean(Kappa_statistic), .groups = "drop"),
    by = join_by(cls)
  ) %>%
  ggplot(aes(S, kappa, label = str_remove_all(cls, "[^A-Z0-9]"))) +
  geom_point(color = "black", pch = 1, size = 3) +
  geom_point(aes(color = cls), pch = 4, size = 3, show.legend = F) +
  ggrepel::geom_label_repel(seed = 7080, family = "mono", fontface = "bold", alpha = 0.5) +
  scale_color_viridis_d(direction = -1) +
  scale_y_continuous(breaks = seq(0, 1, .1)) +
  labs(x = expression(bar(S)), y = expression(bar(kappa))) +
  theme_minimal() +
  theme(
    legend.position = "none",
    panel.border = element_rect(color = "black", linewidth = 1),
    text = element_text(family = "mono")
  )

((p6 / p7) | (p8 / p9)) +
  plot_layout(widths = c(3, 1), guides = "collect") +
  plot_annotation(tag_levels = list(c("A", "C", "B", "D"))) &
  theme(legend.position = "bottom", plot.tag = element_text(size = 16, family = "mono"))
ggsave("Fig2.pdf", dpi = 300)
ggsave("Fig2.png", dpi = 300)

bind_rows(
  mutate(df.eva.nfo, method = "IG"),
  mutate(df.eva.cfs, method = "CFS"),
) %>%
  bind_rows(
    df.zip.con %>%
      separate_wider_delim(attr_key, delim = "_", names = ls.attr.names, cols_remove = F) %>%
      mutate(method = "Freq"),
  ) %>%
  select(method, everything()) %>%
  write_tsv("S5.tsv")

df.csv.con %>%
  pivot_longer(-cls) %>%
  arrange(desc(name)) %>%
  pivot_wider() %>%
  left_join(select(df.csv.tuk, cls, CLD), by = join_by(cls)) %>%
  left_join(summarize(df.zip.sta, S = sprintf("%0.2f (± %.2f)", mean(S), sd(S))), by = join_by(cls)) %>%
  rename(classifier = cls) %>%
  select(classifier, CLD, κ, S, TP, FN, TN, FP, matches("\\.[YN]$", ignore.case = F), κ, everything()) %>%
  arrange(desc(κ)) %>%
  write_tsv("S6.tsv")

df.csv.tuk %>%
  write_tsv("S7.tsv")

df.zip.sta %>%
  write_tsv("S8.tsv")

df.log.mod %>%
  write_tsv("S9.tsv")
```
