## Supplemental code and data. for "Multi-feature Classification to Improve Colorimetric Loop-Mediated Isothermal Amplification Fidelity": README.md

# Manifest

- "data/WEKA-{classifier}-{seed}.out.txt" files are standard output.
- "data/WEKA-{classifier}-{seed}.err.txt" files are standard errer.
- "data/WEKA-{classifier}-{seed}.out.csv" files are classifier output logs.
- "data/WEKA-{classifier}-{seed}.out.zip" files are classifier split logs.
- "data/WEKA-{classifier}-{seed}.log.txt" files are grid search logs.
- BLAST+ query "blast-query.fna".
- BLAST+ hits subset "blast-hits-subset.tsv" file.
- *.{R,py,sh} are experiment scripts
- *.jpg are LAMP assay photos
- *.{pdf,png} are the manuscript figures
- *.txt are time output
- *.pptx has supplemental Figure S1


# Dependencies

- Install [WEKA](https://sourceforge.net/projects/weka/)
- Install [PSET](https://github.com/biolaboro/PSET)
- For the R scripts, install [tidyverse](https://tidyverse.org/), [openxlsx2](https://janmarvin.github.io/openxlsx2/), [patchwork](https://patchwork.data-imaginist.com/), [ggrepel](https://ggrepel.slowkow.com/) and [archive](github.com/r-lib/archive).


# Template search

Run the template search using the provided subset of the BLAST hits as an example. A subset is provided since the original hits table is over 1GB.

- The first R script reads primer data from the supplement and returns a set of subject accession ranges in the "blast-saccver-subset.tsv" file.
- Next, download the subject sequence subsets using NCBI E-utilities.
- The second R script reads primer data from the supplement and results from the first script to demarcate the template with primer locations "blast-definition-subset.fna" file. 

```bash
RScript --vanilla filter1.R
cat blast-saccver-subset.tsv | 
while read -r id start stop; do 
  curl "https://eutils.ncbi.nlm.nih.gov/entrez/eutils/efetch.fcgi?db=nuccore&id=$id&strand=1&seq_start="$start"&seq_stop=$stop&rettype=fasta&retmode=text" 2> /dev/null;
  sleep 2;
done > blast-saccver-subset.fna
RScript --vanilla filter2.R
```


# Feature extraction

The following reads directly from supplement.xlsx and writes S3.csv, S3.tsv, and S3.arff.

```bash
python featurize.py
```


# ML

Run the WEKA Experiment workflow on the command line. 

- Replace "weka-3-9-7/weka.jar" with the location of the WEKA JAR file. 
- Replace 196 with the number of CPUs.
- Make sure to install the MultiSearch plugin beforehand.

```bash
bash WEKA.sh weka-3-9-7/weka.jar S3.arff 196 2>&1 | tee WEKA-run.log.txt
```

On completion, move all of the outputs to a folder called "data" before proceeding.


# Figures and tables

The following plots and writes figures and tables.

```bash
Rscript --vanilla plots.R
```


# LAMP

The "supplement.pptx" contains Figure S1. Raw and edited files are "2026_April29_LAMP_IMG_20260430_054014375_raw.jpg" and "2026_April29_LAMP_IMG_20260430_054014375_edited.jpg" respectively.
