## Supplemental code and data. for "Multi-feature Classification to Improve Colorimetric Loop-Mediated Isothermal Amplification Fidelity": WEKA.sh

```
#!/usr/bin/env bash

# script parameters

jar="${1:-/Applications/weka-3.9.7.app/Contents/app/weka.jar}"
arf="${2:-S3.arff}"
cpu="${3:-8}"


# seeds

seeds=(109 127 157 179 191 211 241 277 283 331)


# analyze features

evaluators=(
    "weka.attributeSelection.CfsSubsetEval"
    "weka.attributeSelection.InfoGainAttributeEval"
)
searchers=(
    "weka.attributeSelection.BestFirst -D 1 -N 5"
    "weka.attributeSelection.Ranker -T -1.7976931348623157E308 -N -1"
)
threads=(
    "1"
    ""
)

for i in "${!evaluators[@]}"; do
    for seed in "${seeds[@]}"; do
        out="WEKA-${evaluators[$i]##*.}-$seed.out.txt"
        err="WEKA-${evaluators[$i]##*.}-$seed.err.txt"
        echo "${evaluators[$i]##*.} $seed..." && \
        java \
            -Xmx8g \
            --add-opens java.base/java.lang=ALL-UNNAMED \
            -cp "$jar" \
            "${evaluators[$i]}" \
            -i "$arf" \
            -s "${searchers[$i]}" \
            -x 10 \
            -n "$seed" \
            -M \
            ${threads[$i]:+-P "$cpu"} \
        1> >(tee "$out") \
        2> >(tee "$err")
    done
done


# optimize classifiers

classifiers=(
    "weka.classifiers.rules.ZeroR"
    "weka.classifiers.rules.OneR"
    "weka.classifiers.trees.DecisionStump"
    "weka.classifiers.bayes.NaiveBayes"
    "weka.classifiers.bayes.BayesNet"
    "weka.classifiers.trees.J48"
    "weka.classifiers.trees.RandomForest"
)
search1="weka.core.setupgenerator.MathParameter -property filter.filters[2].search.numToSelect -min 1.0 -max 20.0 -step 1.0 -base 10.0 -expression I"
search2=(
    ""
    ""
    ""
    ""
    "weka.core.setupgenerator.MathParameter -property classifier.searchAlgorithm.maxNrOfParents -min 1.0 -max 3.0 -step 1.0 -base 10.0 -expression I"
    "weka.core.setupgenerator.MathParameter -property classifier.minNumObj -min 2.0 -max 6.0 -step 2.0 -base 10.0 -expression I"
    "weka.core.setupgenerator.MathParameter -property classifier.maxDepth -min 2.0 -max 4.0 -step 1.0 -base 10.0 -expression I"
)
search3=(
    ""
    ""
    ""
    ""
    "weka.core.setupgenerator.MathParameter -property classifier.estimator.alpha -min 0.1 -max 1.0 -step 0.3 -base 10.0 -expression I"
    "weka.core.setupgenerator.MathParameter -property classifier.confidenceFactor -min 0.125 -max 0.5 -step 0.125 -base 10.0 -expression I"
    "weka.core.setupgenerator.MathParameter -property classifier.bagSizePercent -min 40.0 -max 100.0 -step 20 -base 10.0 -expression I"
)
params=(
    ""
    ""
    ""
    ""
    "-- -D -Q weka.classifiers.bayes.net.search.local.K2 -- -P 1 -S BAYES -E weka.classifiers.bayes.net.estimate.SimpleEstimator -- -A 0.5"
    "-- -C 0.25 -M 2"
    "-- -P 100 -I 100 -num-slots 1 -K 0 -M 1.0 -V 0.001"
)
subseeds=("" "" "" "" "" "" "1")

for i in "${!classifiers[@]}"; do
    for seed in "${seeds[@]}"; do
        cls="${classifiers[$i]##*.}"
        csv="WEKA-$cls-$seed.out.csv"
        zip="WEKA-$cls-$seed.out.zip"
        log="WEKA-$cls-$seed.log.txt"
        out="WEKA-$cls-$seed.out.txt"
        err="WEKA-$cls-$seed.err.txt"
        echo "${classifiers[i]##*.} $seed..." 
        java \
            -Xmx8g \
            --add-opens java.base/java.lang=ALL-UNNAMED \
            -cp "$jar" weka.experiment.Experiment \
            -r \
            -verbose \
            -L "$seed" \
            -U "$seed" \
            -T "$arf" \
            -N "${cls}-${seed}" \
            -D "weka.experiment.CSVResultListener -O $csv"  \
            -P weka.experiment.CrossValidationResultProducer -- \
                -X 10 \
                -D \
                -O "$zip" \
                -W weka.experiment.ClassifierSplitEvaluator -- \
                    -P \
                    -W weka.classifiers.meta.MultiSearch -- \
                        -E KAP \
                        -class-label 1 \
                        -search "$search1" \
                        ${search2[$i]:+-search "${search2[$i]}"} \
                        ${search3[$i]:+-search "${search3[$i]}"} \
                        -algorithm "weka.classifiers.meta.multisearch.DefaultSearch -sample-size 100.0 -initial-folds 5 -subsequent-folds 5 -initial-test-set . -subsequent-test-set . -num-slots $cpu" \
                        -log-file "$log" \
                        -S "$seed" \
                        -W weka.classifiers.meta.FilteredClassifier -- \
                            -F "weka.filters.MultiFilter \
                                -F \"weka.filters.supervised.attribute.Discretize -R first-last -precision 6\" \
                                -F \"weka.filters.supervised.attribute.AttributeSelection -E \\\"weka.attributeSelection.InfoGainAttributeEval\\\" -S \\\"weka.attributeSelection.Ranker -N 100\\\"\" \
                                -F \"weka.filters.supervised.attribute.AttributeSelection -E \\\"weka.attributeSelection.CfsSubsetEval -M\\\" -S \\\"weka.attributeSelection.GreedyStepwise -R -T -1.7976931348623157E308 -N -1\\\"\"" \
                            -S "$seed" \
                            -W "${classifiers[$i]}" \
                                ${params[$i]:+${params[$i]}} \
                                ${subseeds[$i]:+-S "$seed"} \
            1> >(tee "$out") \
            2> >(tee "$err")
    done
done
```
