## Supplementary figures and images for "Multi-feature Classification to Improve Colorimetric Loop-Mediated Isothermal Amplification Fidelity"

### 2026_April29_LAMP_IMG_20260430_054014375_edited.jpg

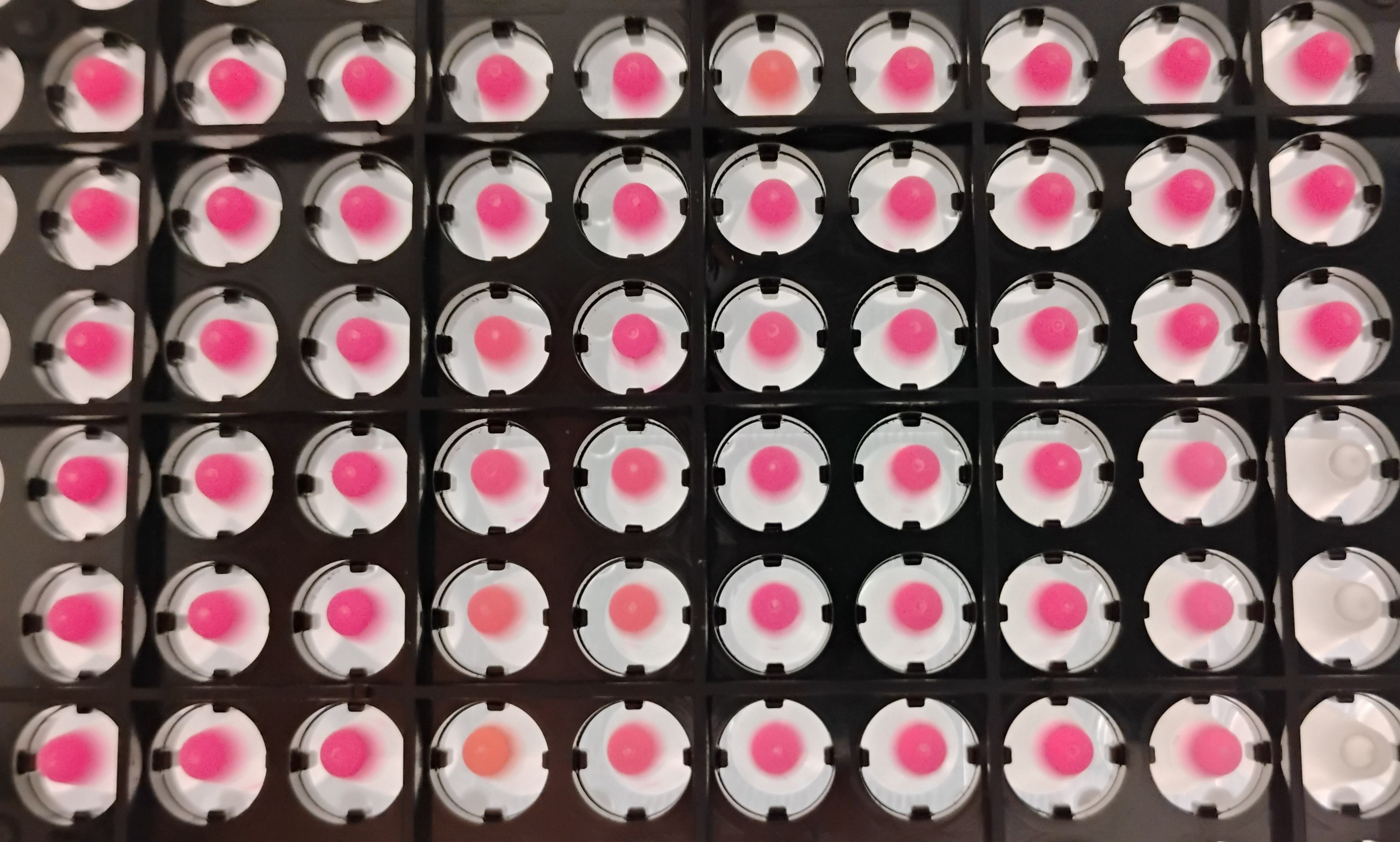

### 2026_April29_LAMP_IMG_20260430_054014375_raw.jpg

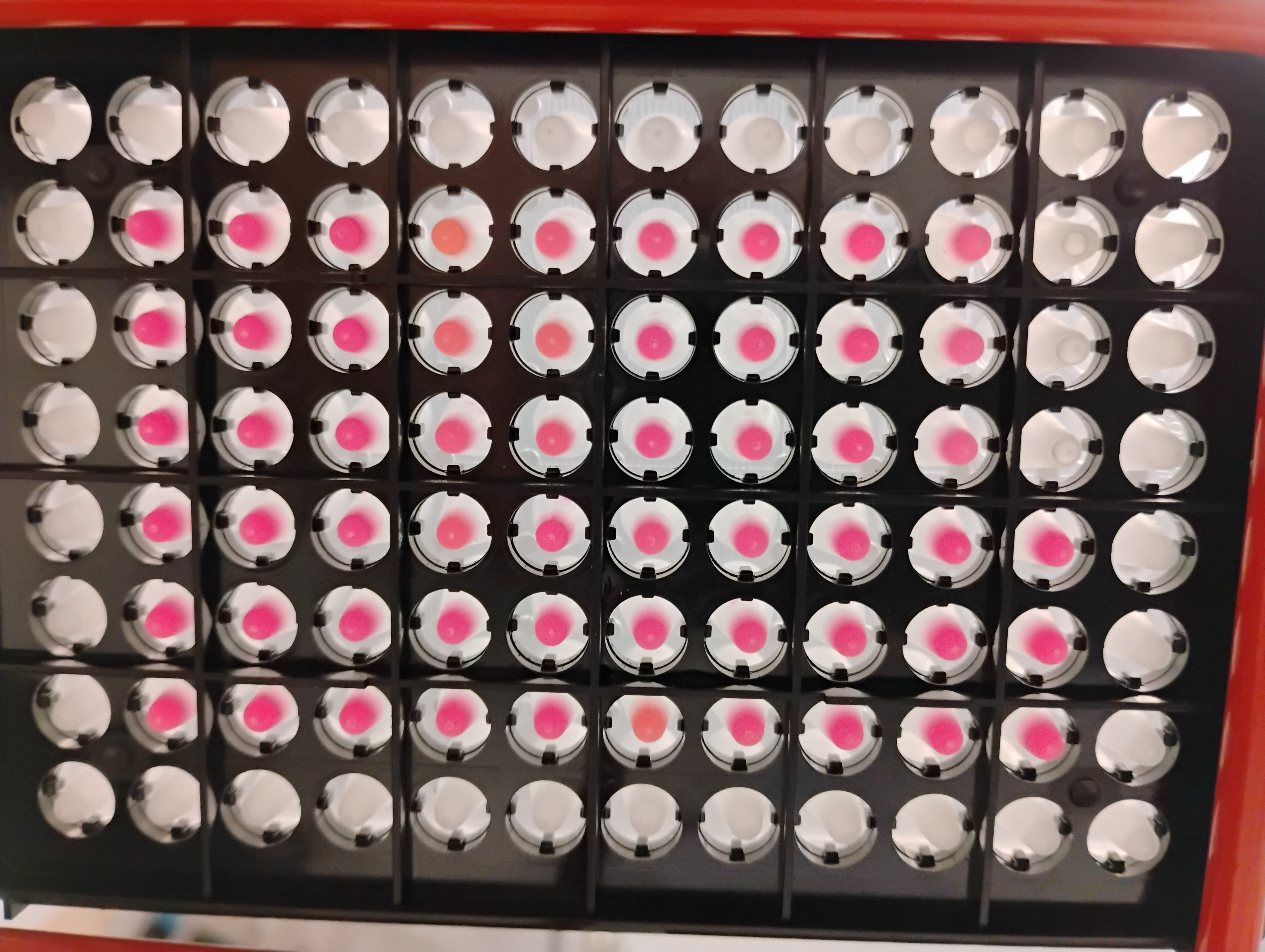

### Fig1.pdf

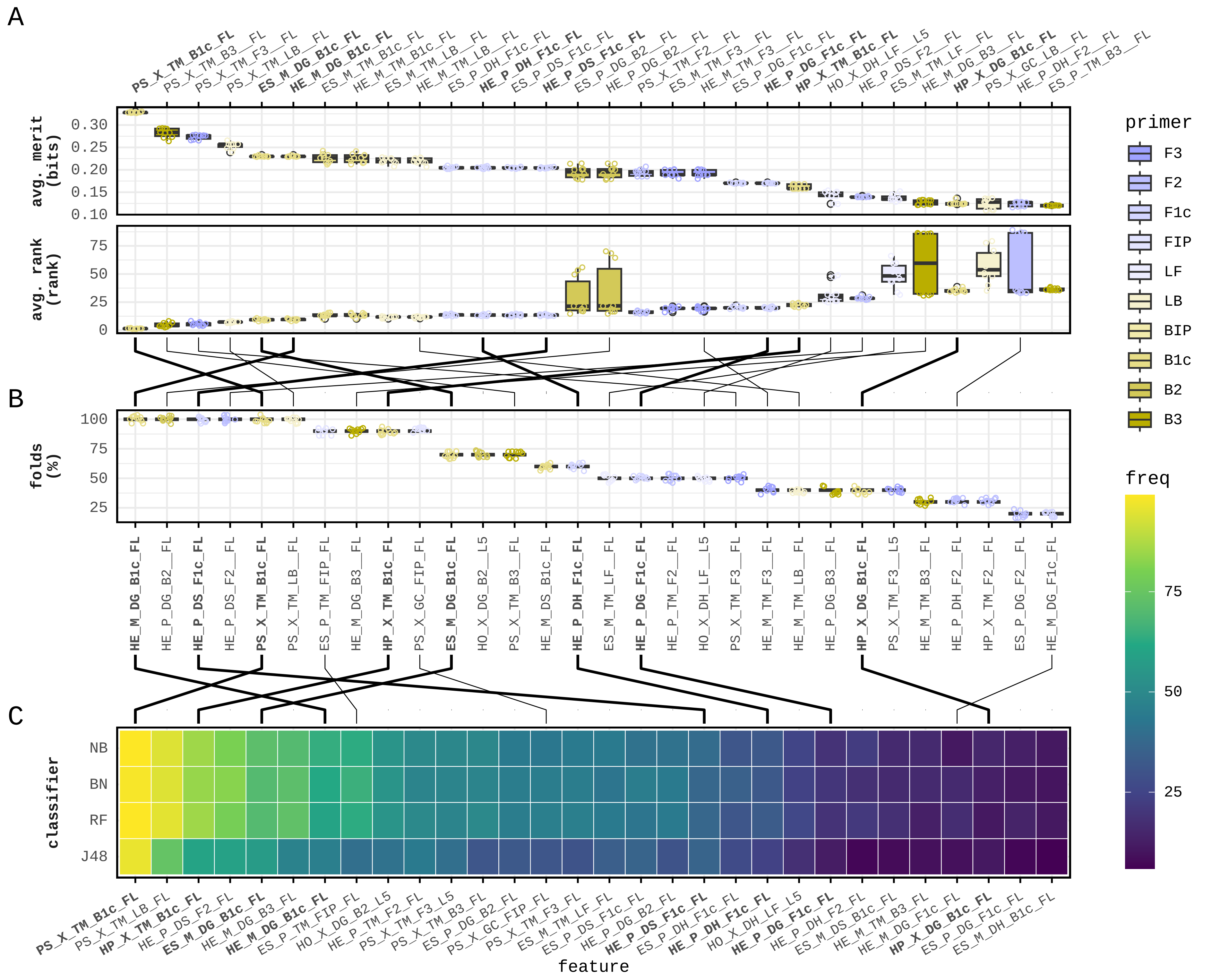

### Fig1.png

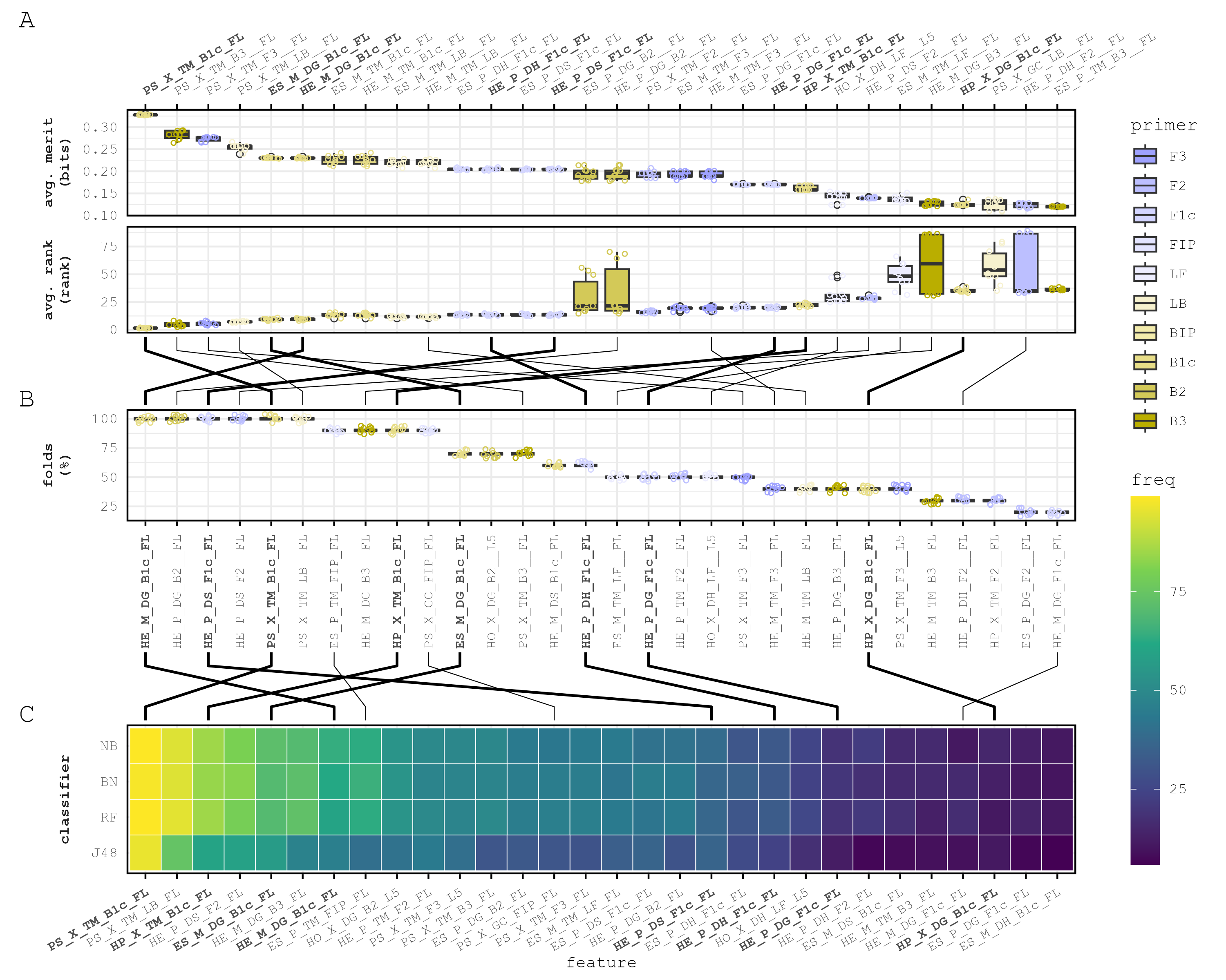

### Fig2.pdf

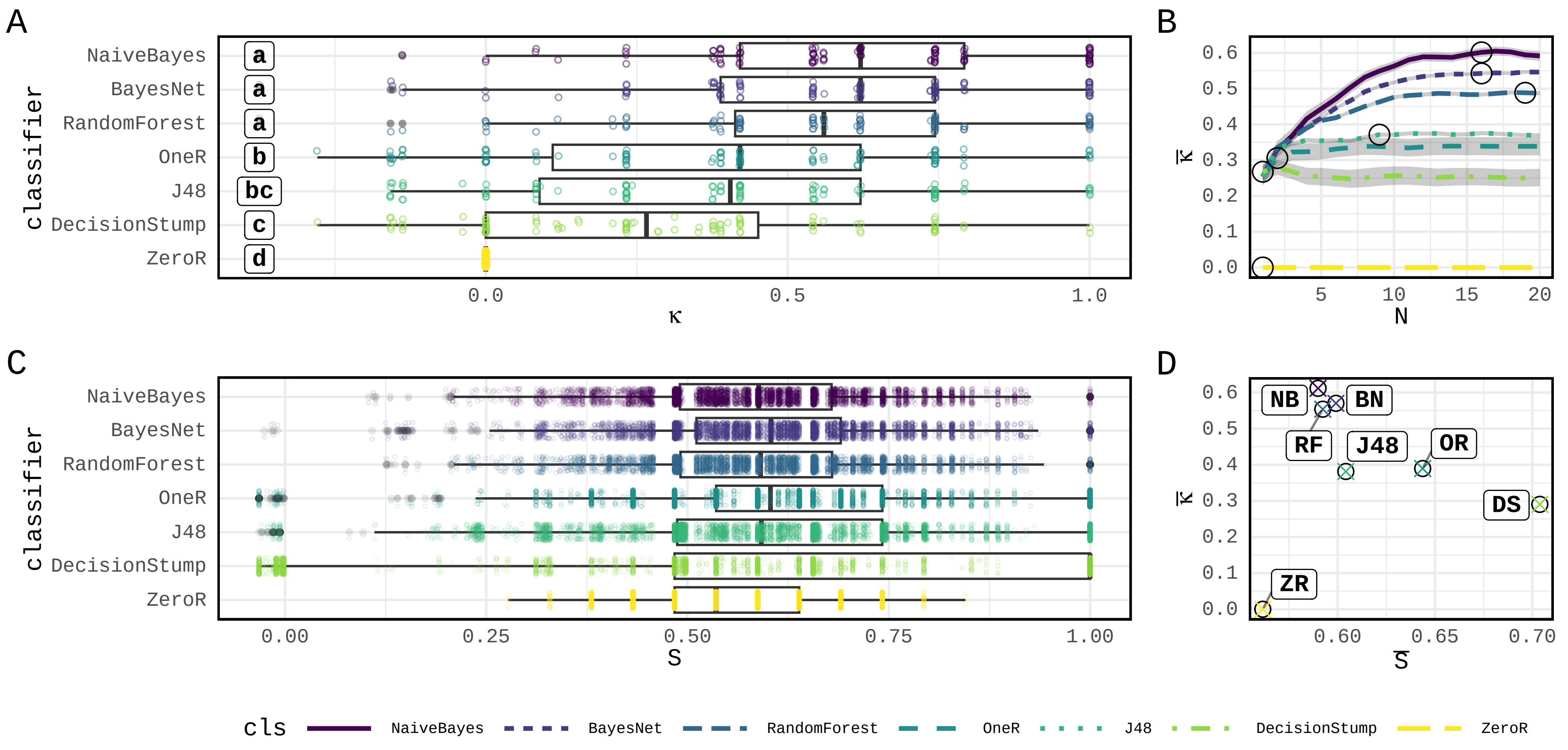

### Fig2.png

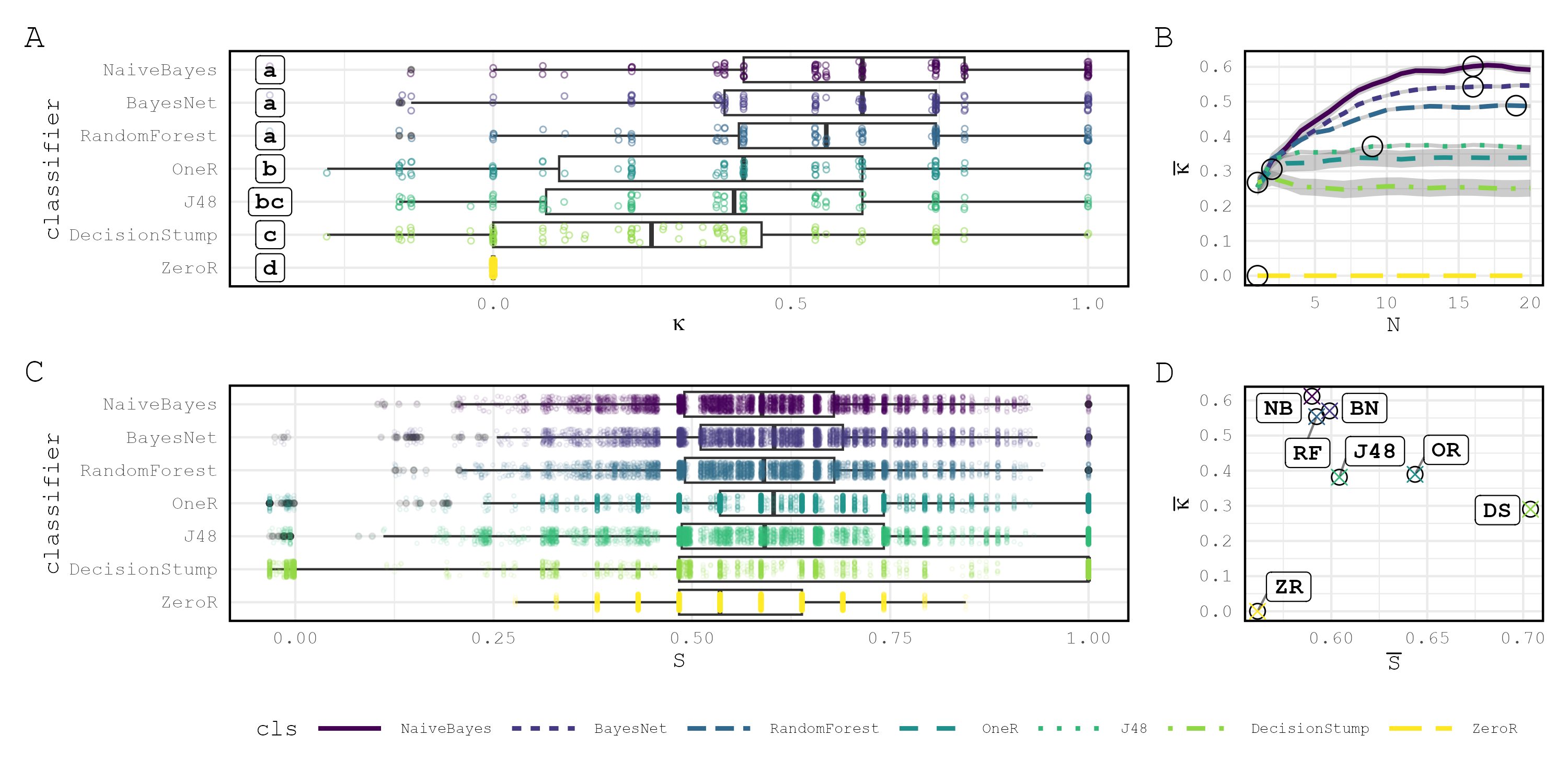
